# Laboratory evolution of the bacterial genome structure through insertion sequence activation

**DOI:** 10.1101/2024.07.14.599650

**Authors:** Yuki Kanai, Atsushi Shibai, Naomi Yokoi, Saburo Tsuru, Chikara Furusawa

**Affiliations:** Department of Biological Sciences, Graduate School of Science, The University of Tokyo, 7-3-1 Hongo, Bunkyo-ku, Tokyo 113-8654, Japan; Center for Biosystem Dynamics Research, RIKEN, 6-2-3 Furuedai, Suita, Osaka, 565-0871, Japan; Universal Biology Institute, Graduate School of Science, The University of Tokyo, 7-3-1 Hongo, Bunkyo-ku, Tokyo 113-8654, Japan

## Abstract

The genome structure critically impacts bacterial physiology, ecology, and evolution. However, its evolution, driven by transposons called insertion sequences (IS), has been challenging to track in laboratories due to its slow pace. Here, we accelerated this process by introducing multiple copies of a high-activity IS into *Escherichia coli*. Mimicking bursts of IS copies in host-restricted endosymbionts and pathogens, we evolved the bacteria under relaxed neutral conditions. Within ten weeks, we observed a median of 24.5 IS insertions per genome, comparable to a decade of wild-type evolution. Long-read sequencing revealed extensive IS-mediated genome rearrangements, resulting in novel IS variants and genome size changes exceeding ±5 %. By achieving such drastic genome evolution under relaxed selection, our study establishes a baseline for assessing the fitness effects of IS insertions, genome size changes, and rearrangements. This work paves the way for experimentally studying bacterial genome structure evolution, complementing analyses of genome structures in nature.

## Introduction

The genome structure, including gene number and arrangement, critically impacts bacterial physiology, ecology, and evolution (Rodríguez-Gijón et al., 2022; Touchon and Rocha, 2016). A key driver of its evolution is the insertion sequence (IS), ubiquitous DNA transposons in bacterial genomes, activating, disrupting, and reordering genes through recombination (Siguier et al., 2014). For instance, most genome rearrangements during the two decades of the long-term evolution experiment of *Escherichia coli* (LTEE) were mediated by ISs (Raeside et al., 2014). In nature, ISs are particularly active in bacterial endosymbionts and pathogens, whose genomes can harbor hundreds of IS copies (McCutcheon and Moran, 2012; Dekker, 2024). The stable and possibly nutrient-rich environments within hosts combined with frequent population size bottlenecks during transmission between hosts relax selection, allowing the accumulation of weakly deleterious mutations including the burst of IS copies (Moran and Plague, 2004). ISs can disrupt genes upon insertion and the proliferated ISs can delete sequences through tandem deletions and homologous recombinations, leading to genome reduction, even down to one-tenth of the size of free-living relatives (Plague et al., 2008).

Despite the abundance of sequence data, inferring the dynamics including the intermediate steps of genome structure evolution can be difficult (Kanai et al., 2022). Moreover, disentangling the potential factors driving these dynamics is fundamentally challenging, as the conditions of evolution in nature cannot be controlled. Laboratory evolution offers an alternative approach, allowing researchers to directly observe evolution under controlled conditions, track the dynamics, and test evolutionary hypotheses (Kawecki et al., 2012), including the role of ISs in genome reduction.

Previous studies struggled to observe IS-mediated evolution in the laboratory primarily due to its slow pace. The model bacterium *E. coli* has a genome that includes almost all the genes of the genome-reduced aphid endosymbiont *Buchnera aphidicola* (Shigenobu et al., 2000) and has pathogenic strains harboring hundreds of ISs (Hawkey et al., 2020). This suggests that *E. coli* has the potential for intense IS-mediated reductive genome evolution in the laboratory. However, IS transposition in *E. coli* typically occurs only once per year (every few thousand generations) (Hawkey et al., 2020; Lee et al., 2016; Consuegra et al., 2021). Plague attempted to simulate natural IS expansions by evolving *E. coli* for 4000 generations under relaxed selection, but failed to detect any IS expansion (Plague et al., 2011). To observe the IS-mediated genome evolution of *E. coli*, previous studies either required decades, as in the LTEE (Consuegra et al., 2021), or involved evolving hundreds of lines in parallel, as in the mutation accumulation (MA) experiment by Foster’s group (Lee et al., 2016). These studies highlight the need for a more efficient method to study IS-mediated genome evolution under well-defined conditions.

Here, we introduce a method to rapidly observe IS-mediated genome evolution in the laboratory. Plague attributed their failure to observe IS expansion to the low IS activity of the wild-type strain of *E. coli* compared to bacteria experiencing bursts of IS activity in nature (Plague et al., 2011). Simulating the conditions of such bursts, we inserted multiple copies of high-activity ISs into the genome of *E. coli* and evolved 44 lines under a stable, nutrient-rich, and low-population size condition.

This approach enabled us to observe extensive IS-mediated genome structure evolution in just ten weeks. Our study provides a reference for studying the fitness effects of IS insertions, genome size changes, and rearrangements and demonstrates a powerful method for experimentally investigating bacterial genome structure evolution under controlled laboratory conditions.

## Results

### The construction of ancestor strains

We designed a derivative of IS*1*, a major IS of *E. coli* (IS*1*-YK2X8; Fig. 1.A). The IS was tailored to induce rapid IS-mediated structural rearrangements in *E. coli* as follows. A high-activity mutant of the transposase gene (*tpn*) (Sekine and Ohtsubo, 1989) was placed under a strong inducible promoter (P_Tet_). The promoter is repressed by the TetR repressor expressed from *tetR* inserted within the IS and is derepressed when anhydrotetracycline (aTc) is supplemented to the growth medium. To facilitate the rapid observation of IS-mediated genome evolution, we introduced multiple copies of IS*1*-YK2X8. This was achieved by introducing a copy of *rfp* in the IS and using red fluorescence as a proxy of IS copy number. After introducing a copy of IS*1*-YK2X8 to an IS-less *E. coli* strain MDS42 (Pósfai et al., 2006), we repeatedly selected for brighter red fluorescence via fluorescence-activated cell sorting (FACS) and colony picking under blue/green LED (Fig. 1.B).

**Fig. 1.**
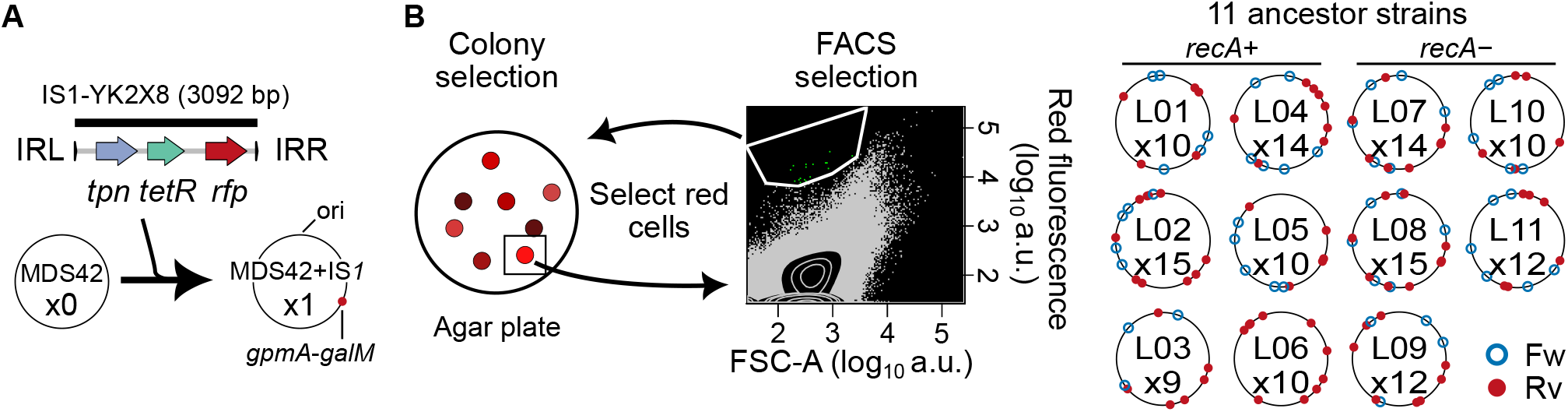
The construction of ancestor strains with multiple copies of high-activity ISs. **(A)** To achieve rapid IS-mediated evolution we constructed *E. coli* strains with high IS activity. First, a copy of an inducible IS*1*-YK2X8 was introduced into the *gpmA*-*galM* locus of IS-less *E. coli* MDS42 by lambda red recombination. *tpn*: transposase gene, *tetR*: tet repressor gene, *rfp*: red fluorescent protein gene, IR: (L: left; R: right) inverted repeat. **(B)** Then, IS was accumulated in the genomes by selecting cells with brighter red fluorescence by fluorescence-activated cell sorting (FACS) and colony picking under blue/green LED. For FACS, bright cells within manually drawn gates (white) were collected. ‘FSC-A’ indicates forward scatter value, used as a proxy for cell size. **(C)** The positions and numbers of IS copies in genomes of the 11 ancestor strains used for the subsequent mutation accumulation experiment. The numbers after ‘x’ indicate the copy numbers. The circular genomes are drawn to scale with the origin of replication (ori) at the top. Fw and Rv: ISs located on the forward and reverse strands, respectively.

We prepared six strains with *recA* and five without *recA* to observe the potential effect of genetic backgrounds on IS-mediated genome evolution (Fig. 1.C). *recA* is the major enzyme in the recombinational repair system in bacteria (Wiktor et al., 2021). The loss of related genes is a characteristic of genome reduction in endosymbionts (McCutcheon and Moran, 2012), and its loss affects the rate of recombination between ISs (Reams et al., 2012). Even so, the effect of *recA* loss on IS-mediated genome evolution was not investigated in Foster’s group’s MA of wild-type *E. coli*, despite running the experiment with knockouts of other repair pathways (Lee et al., 2016).

### Expansion of ISs under relaxed selection

We evolved the strains under conditions simulating the relaxed selection considered to cause IS expansion in nature (Fig. 2.A). When streaked on agar plates, *E. coli* forms colonies derived from single cells. Propagating a single colony every passage reduces the effective population size (*N*_*e*_), allowing the accumulation of weakly deleterious mutations by genetic drift. Laboratory evolution using this method is called MA experiments (Elena and Lenski, 2003). We subjected the IS-accumulated strains to MA (reducing *N*_*e*_), used the nutrient-rich Luria-Bertani medium (allowing more genes to be lost), and achieved high activity of IS by adding aTc to the medium. We evolved four evolutionary lines from every eleven IS-accumulated strains (44 lines). We denote the lines as L01-2: the second line derived from the first ancestor strain, L01. To capture the dynamics of genome evolution, we sequenced genomes at two time-points: post-eight and post-twenty passages.

**Fig. 2.**
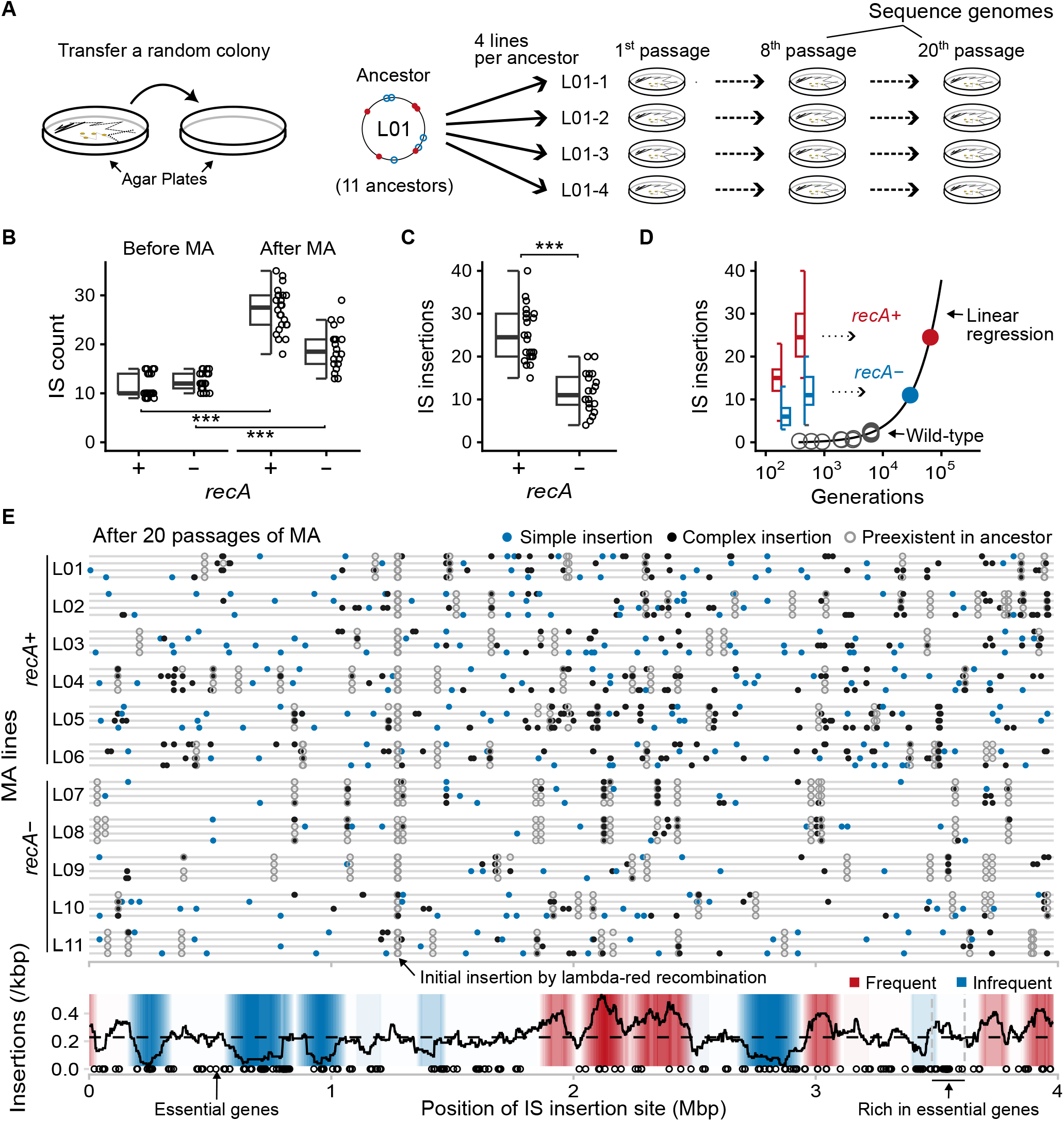
IS expansion after 10 weeks of neutral evolution. **(A)** Overview of the mutation accumulation experiment (MA). Single-cell bottlenecks were imposed on every passage to achieve relaxed neutral evolution. Genomes were sequenced after eight and twenty passages. **(B, C)** IS counts and new insertion sites after MA. *** : *P* = 4.1 × 10^−16^, 7.2 × 10^−6^, 7.1 × 10^−6^ for (B) paired t-test (*recA*^+^: *n* = 24, *recA*^−^: *n* = 20) and (C) Wilcoxon rank-sum test (*n* = 44), respectively. **(D)** Comparison of new insertion site increases with a previous MA of wild-type *E. coli* (Lee et al., 2016). Linear regression (black line) estimates the number of generations required for wild-type MA to reach insertion site counts observed in our study (filled colored circles). Note the x-axis is in log scale. Boxplots show the medians (center line), quartiles (box limits), and 1.5x interquartile ranges (whiskers). **(E)** Distribution of IS insertions after twenty passages of MA. Upper: Positions of IS insertion sites based on the coordinates of MDS42 + IS*1*. Each horizontal line corresponds to one of the 44 evolutionary lines. Blue circles indicate simple insertions, while black circles represent complex insertions (those associated with recombinations). Open circles denote pre-existing ISs before MA. Lower: IS insertion frequency throughout the genome. The line graph represents the mean insertion frequencies within 100 kbp windows sampled every 1 kbp (*n* = 3981). Color depths indicate the number of windows with significant IS insertion accumulation or depletion. Two dashed lines around 3.5 Mbp indicate the essential gene-rich IS-empty zone identified in the LTEE (Consuegra et al., 2021).

After approximately 540 generations (twenty passages, ten weeks), ISs increased by an average of 11.6 copies per genome (Fig. 2.B, Table 1). The *recA*^+^ strains showed an increase of 15.7 (9–23) copies, while the *recA*^−^ strains showed an increase of 6.7 (0–17) copies. ANOVA, with *recA* +/– and the pre-MA IS count as factors, indicated the increase in *recA*^+^ was significantly higher (*F*_1,41_ = 40.0, *P* = 1.5 × 10^−7^). In contrast, the pre-MA IS count had non-significant effects on the increase (*F*_1,41_ = 2.4, *P* = 0.13, linear regression slope = −0.48 (increase/pre-MA count), Supplementary Fig. 3).

**Table 1.**
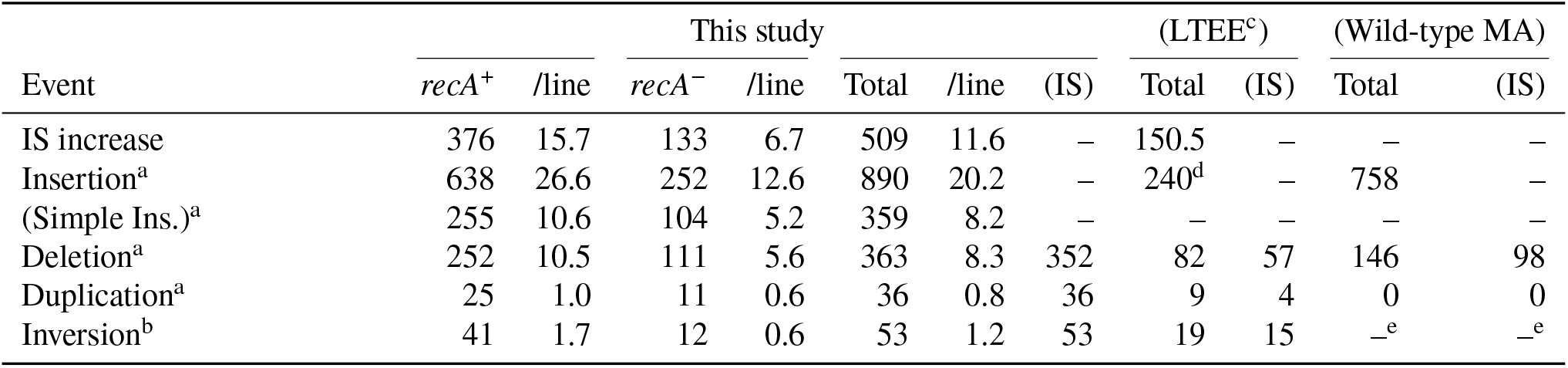
Events during the ten weeks of MA.

To compare the observed copy number increase with the MA of wild-type *E. coli* by Foster’s group without assembled genomes (Lee et al., 2016), we analyzed the number of IS insertion sites: the inserted position of ISs in the coordinates of the IS-excluded genome sequence of a predecessor genome. Comparing the genomes before, eight, and twenty passages, we detected 890 IS insertion sites (Table 1). To highlight the rapid IS expansion observed, we also compared the genomes after twenty passages directly with the ancestor genomes. Remarkably, a median of 24.5 sites per line was detected in *recA*^+^ (Fig. 2.C), a number comparable to roughly 6.5 × 10^4^ generations or 9.0 years of MA using wild-type *E. coli* (Lee et al., 2016) (Fig. 2.D; linear regression, *n* = 15, *R*^2^ = 0.91, *y* = 3.8 × 10^−4^*x* − 0.13).

### The distribution of IS insertions

Mapping the insertion sites to the MDS42 genome revealed a predominantly random global distribution of IS insertions (Fig. 2.E), consistent with Foster’s group’s MA (Lee et al., 2016). Taking 100 kbp sliding windows throughout the genome, some windows exhibited significant IS insertion frequencies (*α* < 0.05 in two-sided binomial test with Benjamini-Hochberg correction for false discovery rate, Fig. 2.E, lower). IS*1* can undertake simple insertions but can also cause recombination upon insertion (Biel and Berg, 1984), which we call complex insertions. Focusing on the simple insertions, none of the windows displayed significance (Two-sided binomial test with BH correction for FDR, *P* ≥ 0.46). An exception was the significantly frequent simple insertions in the terminal 1 Mbp region (115/359, Two-sided binomial test, *P* = 0.011).

We expected ISs to avoid inserting near essential genes (as classified by the PEC database for the wild-type *E. coli* strain MG1655 (Kato and Hashimoto, 2007)). However, analysis of 100 kbp windows revealed only a weak, non-significant anti-correlation between the essential gene count and the frequency of IS insertions (linear regression, *n* = 40, *r* = −0.25, *P* = 0.11, 0.39 ± 0.24 (SE) decrease in insertion counts per essential gene, Supplementary Fig. 4). Furthermore, an IS-empty zone rich in essential genes was identified in the LTEE (Consuegra et al., 2021) (3.480 Mbp to 3.615 Mbp in Fig. 2.E, lower), but the corresponding region in our study did not show a significant depletion of IS insertions (One-sided binomial test, *P* = 0.94). MDS42 has a lower proportion of non-essential genes than the ancestor of the LTEE, strain REL606. We speculated that frequent essential genes in other regions, leading to low insertion frequencies, may have obscured the IS-empty zone in our study. Nevertheless, even comparing the insertion frequency per non-essential sequences, the locus had significantly lower insertion frequency compared to the rest of the genome in the LTEE but not in ours (One-sided binomial test, LTEE: *P* = 0.011, This study: *P* = 0.99).

Following a previous analysis of Foster’s MA (Lee et al., 2016), we analyzed the distance distribution of IS insertions relative to preexisting copies of ISs. This confirmed that new ISs were inserted significantly closer to preexisting ISs than expected by random chance, or “local hopping” of ISs (Lee et al., 2016) (Extended Data Fig. 1, Fisher’s combined probability test, 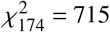, *P* = 1.0 × 10^−66^). However, we noticed that excluding the complex insertions, the distribution of new IS insertion sites was not significantly different from the random distribution (Fisher’s combined probability test, 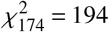, *P* = 0.15). This suggests that the local hopping of ISs was due to complex insertions such as those involving deletions.

### IS-driven genome structure evolution

Given the IS expansion, we next studied the changes in the genome structure (Fig. 3.A, Table 1). We detected a total of 452 large-scale rearrangements, including 53 inversions, 363 deletions, and 36 duplications. 98 % of the rearrangements were associated with ISs, signifying that we successfully observed IS-mediated genome evolution in the laboratory.

**Fig. 3.**
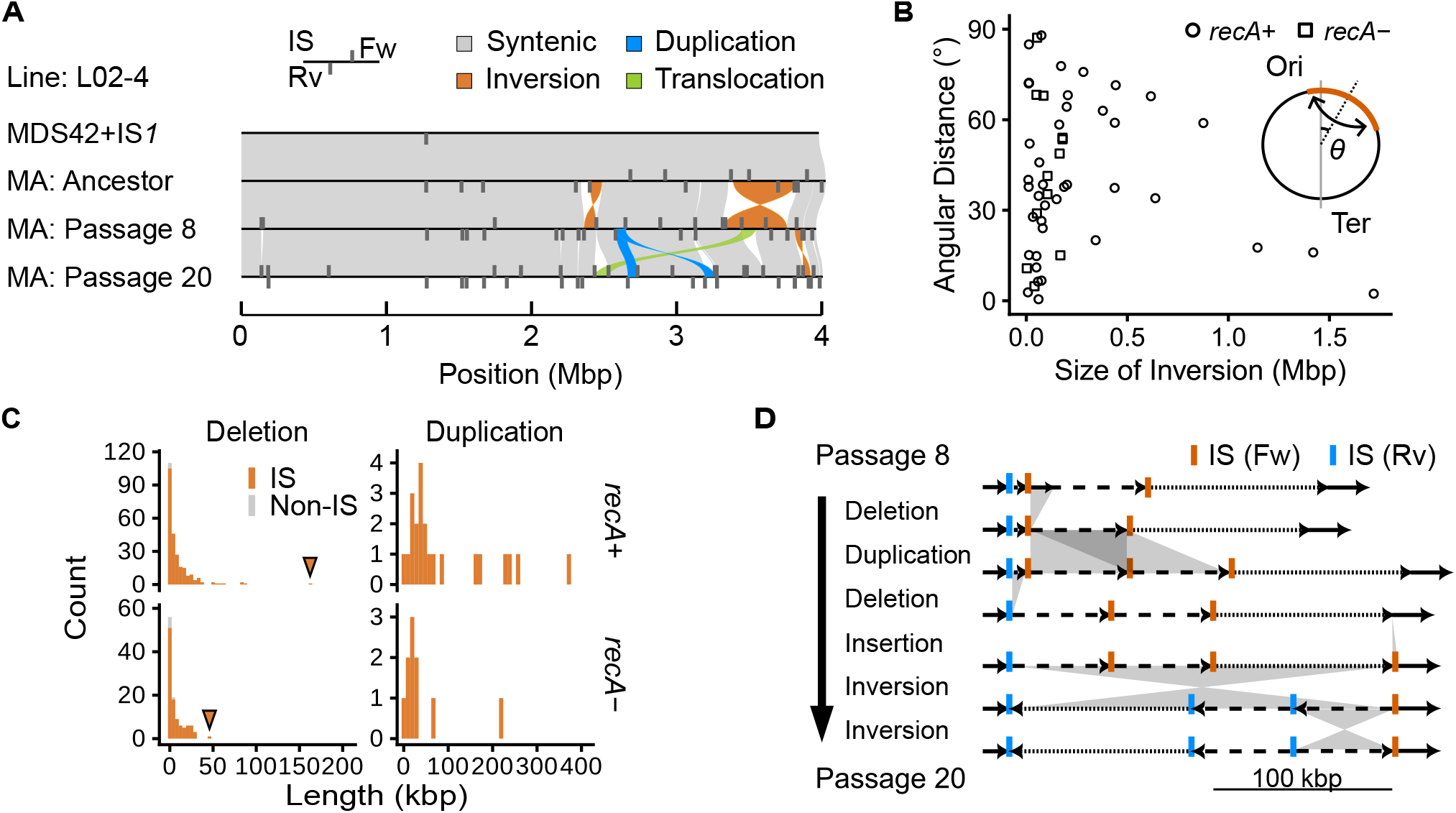
Rearrangements detected after ten weeks of evolution. **(A)** Genome structure evolution of the IS*1*-integrated MDS42 strain, before and after 8 and 20 passages of MA (line L02-4). Genomes are aligned with the origin of replication set at zero. Grey bars represent the positions and orientations of ISs, with the top and bottom bars indicating the direction of the transposase gene. Homologous regions are connected by bands, with colors indicating the type of structural variation identified by SyRI (Goel et al., 2019). Results for other lines are shown in Extended Data Figs. 3–5. **(B)** Inversion sizes and absolute angular distances of inversion centers from the replication origin (*θ*). *θ* = 0 indicates symmetry with respect to the ori-ter axis. **(C)** Length distribution of deletions and duplications. Red indicates mutations with at least one corresponding IS, while gray signifies those that are not. Triangles indicate the largest deletions. **(D)** Hypothetical route of a complex rearrangement observed in L05-4 between passages 8 and 20. Line types represent distinct sequences within the genome. The depicted events represent one of the most parsimonious evolutionary routes inferred from the alignment of the two genomes.

The median size of inversions was 85 kbp (range: 2 kbp– 1.7 Mbp, Fig. 3.B). All three inversions exceeding 1 Mbp were nearly symmetric to the ori-ter axis, consistent with observations of large inversions in nature (Darling et al., 2008).

Deletions ranged from 102 bp to 163 kbp (Fig. 3.C), with the largest deletion being a deletion of a duplicated region (L04-3, Extended Data Fig. 3). Despite the general randomness of IS insertions, the observed deletion sizes were much shorter than expected. Assuming a uniform distribution of deletions from preexisting ISs to random loci that do not contain any essential genes in between, the expected median deletion size was 26 kbp (bootstrap 95 % confidence interval: 24–27 kbp), but the actual median deletion size was only 2516 bp. Nevertheless, the large number of deletions resulted in deletions spanning 1.3 Mbp in total, corresponding to 34 % of the genome (Extended Data Fig. 2). As expected, no essential genes in the PEC and Keio datasets were deleted (Kato and Hashimoto, 2007).

Duplications had a median size of 38 kbp (range: 560 bp–372 kbp, Fig. 3.C). Surprisingly, the majority of the duplications resulted from transpositions of composite transposons with a copy of IS at each end (27/36), rather than tandem duplications.

Inversions and deletions can complicate the detection of IS losses. Nevertheless, there were 25 cases of IS loss without such mutations. The number of losses detected between the eighth and twentieth passages was significantly higher than those up to the eighth passage (20/25, One-sided binomial test, *P* = 0.026). Interestingly, among these 20 losses, 17 were ISs that were newly inserted by simple insertions during the MA up to the eighth passage, rather than ISs inserted before the MA. This proportion was significantly higher than expected based on the fraction of ISs in the genomes that were newly inserted by simple insertions during the MA (One-sided binomial test, *P* = 4.0 × 10^−6^).

The interplay of IS-related mutations drove extensive genome rearrangements through insertions, inversions, deletions, and duplications. Line L05-4 exemplifies this rapid genomic change, displaying a complex combination of these mutations (Fig. 3.D). Despite the complexity, our frequent sequencing ensured that unresolved complex rearrangements were rare and unlikely to have significantly affected the overall mutation statistics presented.

While the rearrangements led to genome size changes ranging from −4.4 % to 9.2 % (Fig. 4.A), overall, deletions were offset by the genome size increase from duplications and IS insertions, resulting in negligible changes in the median genome sizes (*recA*^+^: 1.3 × 10^−3^ %, *recA*^−^: −0.21 %). To account for the low non-essential gene content of MDS42 compared to the ancestor of LTEE and the larger size of ISs used in our study, we adjusted the sizes of deletions based on the non-essential gene content of the parent strains used in the two studies (MA ancestors vs. REL606), and the sizes of IS insertions were reduced to that of the wild-type IS*1*. In addition, for the LTEE, we focused on the non-point-mutator strains, as the point-mutator strains had decelerated the pace of IS insertions (Consuegra et al., 2021). We found that the two studies had a similar number of both median IS insertions (*recA*^+^: 24.5, LTEE: 23.8) and median genome size reduction (*recA*^*+*^: −0.96 %, *n* = 24, bootstrap 95 % confidence interval: (−1.7, 1.2); LTEE: −0.82 %, *n* = 6, bootstrap 95 % confidence interval: (−1.3, −0.035); Fig. 4.B). Interestingly, while the genome size changes strongly correlated with the number of IS insertions in the LTEE (linear regression, *R*^2^ = 0.85, *P* = 0.0033, *n* = 6), our study showed a large variation in genome size changes even among strains with similar numbers of IS insertions, obscuring the correlation (linear regression, *R*^2^ = 0.020, *P* = 0.51, *n* = 24). The largest contributor to the variance in genome size changes in our study was the length of duplications, as indicated by ANOVA among the factors considered: deletions (*F*_1,20_ = 1.6 × 10^6^), duplications (*F*_1,20_ = 7.9 × 10^6^), and IS insertions (*F*_1,20_ = 5.4 × 10^3^) (*n* = 24).

**Fig. 4.**
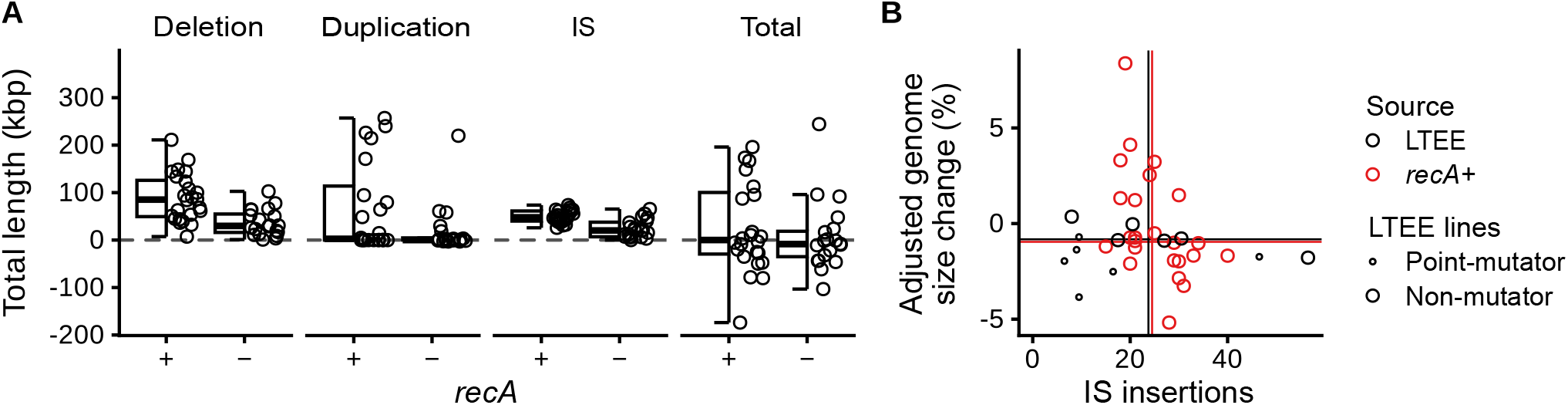
Genome size changes after ten weeks of evolution. **(A)** Total length changes due to the structural mutations. Each data point signifies the overall length variation for a particular type of mutation in an evolutionary lineage. *IS* shows the total length of ISs, while *Total* shows the overall genome size changes. Boxplots show the medians (center line), quartiles (box limits), and 1.5x interquartile ranges (whiskers). **(B)** Genome size changes after adjustments to compare with the LTEE. Lines indicate medians for *recA*^+^ strains in this study and non-mutator strains in the LTEE (Tenaillon et al., 2016).

### The evolution of gene order within the IS

Unexpectedly, various structural variants of ISs were detected (Fig. 5.A). While some of the IS variants were present in the ancestor genomes (L01, L02, L03, L09, and L10), new IS variants emerged in 10/20 lines with pre-existing IS variants and 12/24 lines without pre-existing IS variants. Various mechanisms formed the variants. For instance, L10-3.3.0 (underlined in Fig. 5.A) likely formed through a deletion of the sequence between two ISs, L10-3.2.0 and L10-3.2.1. Other IS variants were formed through the insertion of one IS within another (Fig. 5.B). This arrangement brings homologous sequences into proximity, potentially leading to frequent homologous recombination within an IS, resulting in the further formation of variants (e.g., L10-2.2.20, dashed underlined in Fig. 5.A). Furthermore, the transposition of subsequences from within the nested ISs led to the formation of additional variants (Fig. 5.B).

**Fig. 5.**
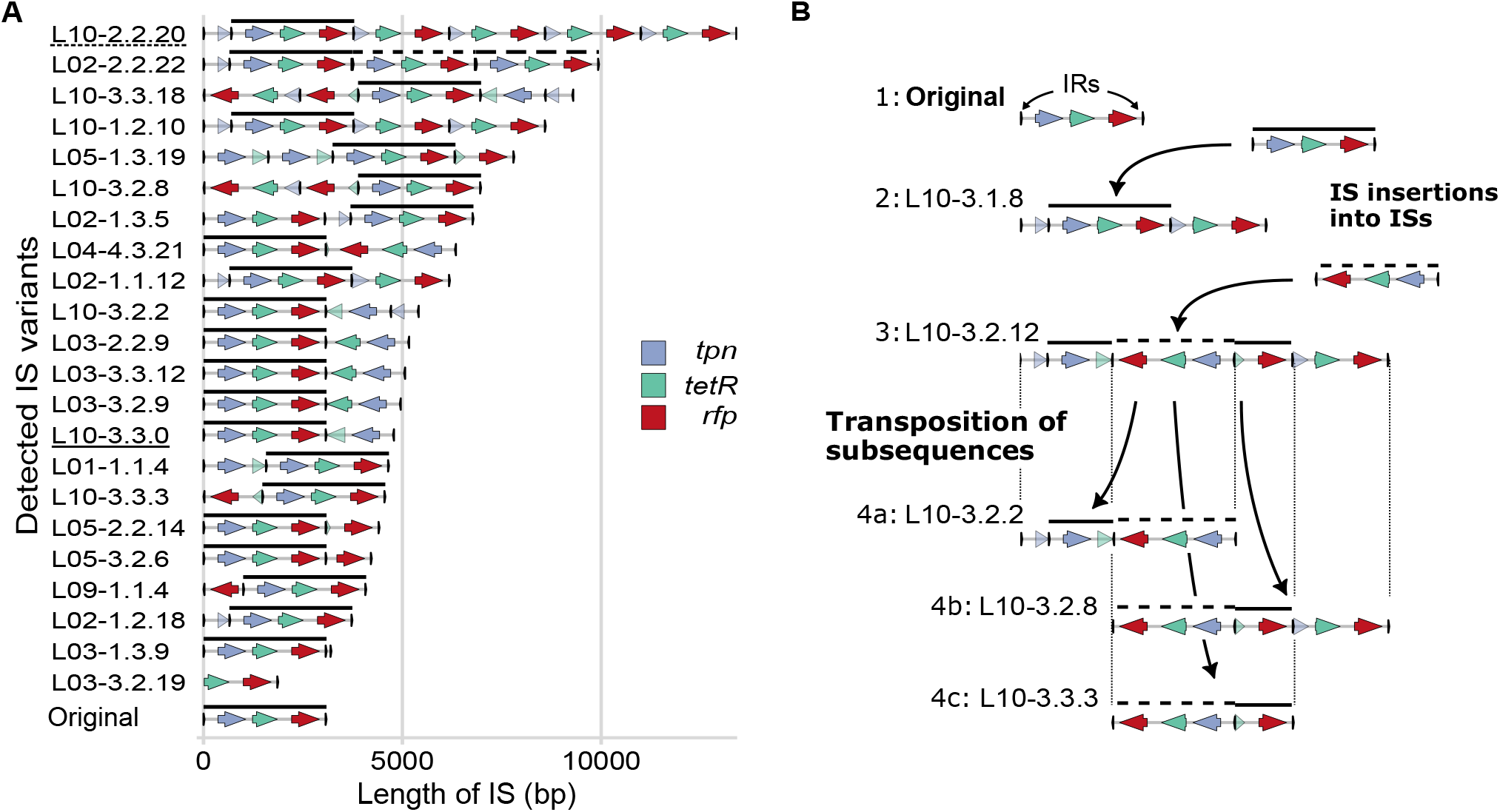
Representative IS variants detected after ten weeks of evolution. **(A)** The representative variants. Each IS is uniquely labeled, e.g., L01-2.3.4 represents the 4th IS in the genome of the 3rd DNA sequencing (1st: Before MA, 2nd: passage 8, 3rd: passage 20) of line L01-2. Full-length ISs are indicated by the lines above genes. Disrupted genes are shown in translucent colors. **(B)** Example of IS variant formation. Steps 1–4 represent a hypothetical scenario inferred from the structures of ISs. The lines above genes distinguish sequences derived from inserted ISs from those from the original IS.

## Discussion

Bacterial genomes are dynamic and can undergo drastic changes. However, two key aspects of genome evolution remain elusive despite accumulating sequence data. First, intermediate evolutionary steps are often obscured, as we previously suggested for the IS-mediated evolution of operons (Kanai et al., 2022). Second, the specific conditions driving evolutionary phenomena are difficult to identify in complex and varying natural environments. Laboratory evolution allows researchers to disentangle such intricacies through controlled experiments (Kawecki et al., 2012) but has been limited by the typically slow pace of genome structure evolution (Plague et al., 2011; Raeside et al., 2014; Lee et al., 2016).

To address these challenges, we developed an *E. coli* strain with high IS activity (Fig. 1) and demonstrated rapid IS-mediated genome evolution under relaxed selection in the laboratory, simulating the natural evolution of symbionts and pathogens. In just ten weeks, we observed numerous IS insertions, IS-mediated duplications, and deletions (Fig. 2, Fig. 3), contributing to over ±5 % genome size changes (Fig. 4). This suggests that, given the right conditions, bacterial genomes can rapidly evolve their structure in a relatively short time, similar to pathogens that rapidly adapt to new hosts and antibiotics (Armbruster et al., 2021; He et al., 2016).

Long-read sequencing enabled us to reveal numerous IS insertions and IS-mediated structural mutations (Table 1). IS variants were also observed (Fig. 5), showcasing our method’s potential to study both the evolution of bacterial genomes and the underexplored evolution of ISs themselves (Kanai et al., 2023). These variants likely represent intermediate stages in the evolution of composite transposons, key elements in spreading antibiotic resistance and pathogenicity (Vandecraen et al., 2017).

By tracking evolution under controlled conditions with accelerated timescales, our approach allows the investigation of fundamental questions surrounding bacterial genome evolution. Our experiment fills the gap of the lack of experimental studies demonstrating extensive genome evolution under relaxed selection and comparing our study with previous studies clarifies notions that previously were only vaguely suggested.

For instance, our experiment supports the notion that bacterial genomes are biased toward deletions (Moran, 2002; Kuo et al., 2009; Sela et al., 2016). In the laboratory, the LTEE showed a decrease in genome sizes (Tenaillon et al., 2016), but it has remained unclear whether the decrease may reflect selective forces or mutation biases. Despite the effective population size of our study (*N*_*e*_ ∼ 10) being much smaller than that of the LTEE (*N*_*e*_ = 3.3 × 10^7^) (Lenski et al., 1991), the decrease in genome sizes in our study and the LTEE were remarkably similar (Fig. 4.B). This supports that the genome reduction in the LTEE reflects genetic hitchhiking of deletions with beneficial mutations (Raeside et al., 2014; Tenaillon et al., 2016).

Notably, duplications that were totally missed in Foster’s MA (Lee et al., 2016) were the major contributors to genome size variation in our study. Since the genomes gained sizes through duplication by composite transpositions (Fig. 3.C), we speculate that the use of wild-type *E. coli* in the previous study might have limited the observation of duplications. Efficient composite transposition requires pairs of ISs to be near each other (Morisato et al., 1983). Whereas the wild-type *E. coli* has only six copies of IS*1* (Lee et al., 2016), with hundreds of kilobase pairs apart, our ancestral strains had over ten copies of IS*1*-YK2X8 (Fig. 1.B), which might have enabled the observation of multiple duplications. Our results suggest that, at least transiently, ISs may not only act as drivers of genome reduction (Moran, 2002) but might also act to increase genome sizes as in other mobile elements (Siozios et al., 2023).

Our study clarifies the mechanisms underlying the coexistence of ISs and bacteria in genomes, which is maintained through a balance of horizontal transfer, transposition, loss, and the fitness effects of IS insertions. Comparative genomic analyses of ISs in natural populations have suggested that ISs are under neutral selection and maintained through transient bursts of transposition (Bichsel et al., 2013; Iranzo et al., 2014). While it was unclear whether ISs could expand in the genome without selection for beneficial IS insertions, our study provides direct evidence that they can (Fig. 2.B). However, while the comparative studies claimed weak purifying selection against IS insertions (Bichsel et al., 2013; Iranzo et al., 2014), the LTEE observed an essential-gene-rich IS-empty zone, suggesting stronger purifying selection (Consuegra et al., 2021). The absence of the IS-empty zone in our study with a smaller effective population size supports this (Fig. 2.E). The previous comparative studies might have underestimated purifying selection against IS insertions by using simplified theoretical models that do not consider the mixed selection pressures of beneficial and deleterious IS insertions, instead only considering their average effects (Bichsel et al., 2013; Iranzo et al., 2014).

Some loci seem to have had a higher frequency of IS-related mutations than others, as newly inserted ISs were more likely to be lost than older ones, and we observed a locus with multiple structural mutations in a short period (Fig. 3.D). We speculate that, in addition to the frequency of composite transpositions, the high ratio of deletions vs. insertions in our study compared to wild-type (41 % vs. 19 %; Table 1) may reflect the different IS positions in the ancestors. Strains with high IS activity or strains with recent IS expansions might better represent IS-mediated genome evolution in nature than wild-type *E. coli* often used.

Through comparing *recA*± strains, we unexpectedly observed reduced IS activity in *recA*-deficient strains (Fig. 2.B), possibly due to the inability to repair double-strand breaks triggered by IS transposase (Reams et al., 2012). Genome-reduced endosymbiotic bacteria often lack recombination genes (McCutcheon and Moran, 2012). While this is commonly thought to promote genome degradation (Dale et al., 2003), our findings suggest it might also lower IS activities, potentially preventing deleterious genome rearrangements.

There are three limitations of note. First, our proposed method requires improvement, as the observed genome evolution remained minor compared to natural evolution, which can involve megabase-pair changes in genome sizes and hundreds of IS copies (McCutcheon and Moran, 2012). Second, to accelerate IS-mediated genome evolution, we modified the regulation of IS activity (Nagy and Chandler, 2004). Identifying the condition triggering IS expansion in nature is necessary to determine the relevance of our results to the early stages of host restriction we aimed to simulate. Thirdly, due to the rapid and extensive structural changes in the genome, we failed to accurately identify some mutations. Sequencing more frequently and improving the accuracy and length of long-read sequencing would help detect complex rearrangements and IS variants.

The slow pace of IS-mediated evolution has been a bottleneck in studying genome evolution, leading to the reliance on comparative studies. Organisms with high IS activity are often slow growers (Miller et al., 2021) or pathogenic (Hawkey et al., 2020). Artificial genome rearrangement methods, such as protoplast fusion (Patnaik et al., 2002; Zhang et al., 2002) or site-specific recombination on synthesized genomes (Dymond et al., 2011) do not reflect typical natural processes. Applying our method to rapidly growing, safe, and experimentally tractable organisms like *E. coli* could serve as a proxy for studying the evolutionary dynamics of unculturable or pathogenic species. We believe our accessible method to observe genome evolution under controlled laboratory conditions stimulates further studies toward an experimentally grounded understanding of the principles behind bacterial genome evolution.

## Methods

### Reagents, strains, and plasmids

We used the following reagents and instruments:

For cell cultures during the MA, we used: LB Broth, Miller (NACALAI TESQUE, Japan, 20068-75), LB Agar, Miller (NACALAI TESQUE, Japan, 20069-65), Anhydrotetracycline hydrochloride (Sigma-Aldrich, USA, 37919), Chloramphenicol (Wako, Japan, 034-10572), Dimethyl Sulfoxide (Wako, Japan, 043-07216), Ethanol (99.5) (Wako, Japan, 057-00456).

The master stock of aTc was prepared at 5 mg/mL in dimethyl sulfoxide, and the working stock was prepared by diluting to 100 µM with ethanol and stored at −20 °C. Chloramphenicol was dissolved in ethanol and stored at −20 °C.

For handling nucleic acids and DNA sequencing, we used: KOD One PCR Master Mix Blue (Toyobo, Japan, KMM-201), Exonuclease V (RecBCD) (New England Biolabs, USA, M0345S), Rapid Barcoding Kit 96 (Oxford Nanopore Technologies, UK, SQK-RBK110.96), Native Barcoding Kit 24 V14 (Oxford Nanopore Technologies, UK, SQK-NBD114), In-Fusion Snap Assembly Master Mix (Takara Bio, Japan, 638948), Phenol/Chloroform/Isoamyl alcohol (25:24:1) (NIPPON Genetics, Japan, 311-90151), Wizard HMW DNA Extraction Kit (Promega, USA, A2920).

We used the following instruments: Infinite F200 multimode plate reader (Tecan, Switzerland), Blue/green LED transilluminator (NIPPON Genetics, Japan, LB-16BG). FACSAria III (BD, USA), Flongle Flow Cell R9.4.1 (Oxford Nanopore Technologies, UK, FLO-FLG001), Minion Flow Cell R9.4.1 (Oxford Nanopore Technologies, UK, FLO-MIN106D), MinION Mk1B (Oxford Nanopore Technologies, UK, MIN-101B), Minion Flow Cell R10.4.1 (Oxford Nanopore Technologies, UK, FLO-MIN114).

We used the following strains and plasmids: *E. coli* MDS42 (Pósfai et al., 2006), BW25113 *recA* (JW2669-KC, NBRP-E.coli at NIG, Japan) (Baba et al., 2006), pKD46 (Datsenko and Wanner, 2000), pCP20 (Cherepanov and Wackernagel, 1995), pAJM.011 (Meyer et al., 2019).

The following plasmids were constructed in this study: pKD46_tetR, pYK2X8 (plasmid with IS*1*-YK2X8). The maps are provided in Supplementary Fig. 1, and sequences will be provided as in the *Data availability*.

### Preparation of strains and plasmids

We designed IS*1*-YK2X8 based on IS*1*, one of the most active IS in wild-type *E. coli* (Lee et al., 2016), to observe IS-mediated evolution in the laboratory. The genetic map of IS*1*-YK2X8 is provided in Supplementary Fig. 1.

To increase IS activity, we made the following modifications to IS*1*. The wild-type transposase gene of IS*1* (*tpn*) has a frameshift in its coding sequence, significantly reducing its activity (Sekine and Ohtsubo, 1989). The A_6_C mutation was introduced to the transposase gene to fix the frameshift to recover IS activity (Sekine and Ohtsubo, 1989). *tpn* was expressed from a strong inducible promoter based on P_LtetO-1_(Kanai et al., 2022). *tetR* from pAJM.011 repressed the promoter (Meyer et al., 2019), preventing unintended IS activity. Anhydrotetracycline (aTc) was added to culture media to cancel the repression and induce the expression of *tpn* to promote IS activity. Starting from an increased copy number of IS would likely increase the rate of IS-mediated genome evolution. To increase the copy number of IS based on fluorescence intensity, we introduced *rfp* (mScarlet-I) from pYK-1N5 (Kanai et al., 2022). Furthermore, we speculated that transcription at the ends of the IS would interfere with the transposase binding to the ends of the IS. To avoid promoter activities, we inserted strong terminators at the ends (*rrnB* T1 and L3S3P21 (Chen et al., 2013)). Inadvertent introduction of promoters was avoided by checking for potential promoter sequences computationally (LaFleur et al., 2022). To avoid recombination within the IS*1* as much as possible, we checked for potentially unstable repetitive sequences using the EFM calculator (Jack et al., 2015). For example, *tpn* overlaps with the terminal sequence of ISs called the inverted repeat (IR). The recombination between the IR and *tpn* may result in the unwanted loss or duplication of *tetR* and *rfp*. We reassigned the codon of the 3^′^-end of *tpn* to remove this homology.

A copy of IS*1*-YK2X8 was introduced into the genome of an IS-free strain of *E. coli*, MDS42 (Pósfai et al., 2006). This strain was used to prevent native ISs from interfering with the IS*1*-YK2X8 activity. The IS was introduced using lambda red recombination (Datsenko and Wanner, 2000) with plasmid pKD46_tetR (Supplementary Fig. 1.A). The plasmid was synthesized by In-Fusion reaction, integrating the *tetR* cassette from pAJM.011 (Meyer et al., 2019) into pKD46 (Datsenko and Wanner, 2000). pKD46_tetR was used instead of the typical pKD46 (Datsenko and Wanner, 2000) to enhance the efficiency of lambda red recombination by repressing leaky IS*1* transposase expression, which may kill cells with IS*1* integrated. The recombination was performed as follows. The cassette with approximately 500 bp overhang, homologous to the *gpmA* and *galM* genes of MDS42, was prepared on a plasmid (pYK2X8, Supplementary Fig. 1.B). The homologous sequences are adjacent in the MDS42 genome, and no sequence is deleted upon integration. The PCR primers to amplify the *gpmA*-IS-*galM* cassette can also amplify a shorter sequence between the *gpmA* and *galM* genes from the genome. To avoid the amplification of genomic *gpmA* and *galM* that do not contain the IS, the plasmid was cured of residual genome DNA pre-PCR by Exonuclease V treatment that deletes linear DNA for thirty minutes following the manufacturer’s protocol. Approximately 1 µg of PCR product was transformed into 80 µL of electrocompetent cells prepared from log-phase MDS42 cells cultured in Super Optimal Broth (SOB) with 0.1 % (w/v) arabinose. After post-culture at 37 °C in SOB with catabolite repression (SOC) medium, cells were plated on LB agar plates with 12.5 µg/mL chloramphenicol at 42 °C to cure pKD46_tetR. IS insertion was confirmed by PCR using primers outside the homologous sequences and the loss of the plasmid was confirmed by the loss of ampicillin resistance.

IS*1* has higher activities at lower temperatures (Reif and Saedler, 1975). Therefore, in the subsequent experiments, cells were cultured at 32 °C when inducing IS activity and at 37 °C otherwise to avoid unintended IS activity.

### IS accumulation using fluorescence as a proxy for IS copy number

Using fluorescence as a proxy for IS copy number, we prepared IS-accumulated strains with two genetic backgrounds: *recA*^+^ and *recA*^−^. *recA*^+^ strains were prepared through cycles of FACS-based selection and colony-based selections (Fig. 1.B).

First, we selected cells by FACS. Four colonies obtained through the genome integration described in the previous section were transferred to LB medium with 12.5 µg/mL chloramphenicol and 100 nM aTc, and incubated at 32 °C with continuous shaking for one overnight. The antibiotic chloramphenicol was added to the media to avoid contamination of cultures by other bacteria. Approximately 100 cells exhibiting the brightest approximately 0.01 % fluorescence were isolated using FACS into 200 µL phosphate-buffered saline (PBS) and cultivated on LB agar plates at 37 °C for two overnights. Twelve bright colonies under a blue/green transilluminator were picked and cultured in a rich defined medium based on standard M9 (Supplementary Table 2) and were considered independent evolutionary lines for the subsequent steps of IS accumulation.

Subsequent selection consisted of FACS-based and colony-based phases. Colonies were diluted in the modified M9 medium with varying concentrations of aTc (0, 1, 10, 100 nM) and incubated at 32 °C for two overnights. The cell cultures with the strongest induction without growth defects were diluted into PBS, and the 0.01 % brightest cells were collected by FACS into 200 µL PBS. The cell mixes were spread onto LB agar plates and incubated at 37 °C to obtain isolated colonies. The three brightest colonies were identified using a custom ImageJ script on photographs taken under a blue/green LED transilluminator, mixed into the modified M9 medium, and used for subsequent rounds of selections. A total of four rounds of FACS-based IS accumulation were performed.

*recA*^−^ strains were prepared similarly. *recA* was deleted before IS accumulation from MDS42 by integrating the *kanR* cassette, which was amplified from the *recA*^−^ strain of the Keio collection (Baba et al., 2006), using lambda red recombination (Datsenko and Wanner, 2000). The *kanR* cassette was then removed by FLP recombinase expressed from pCP20 (Cherepanov and Wackernagel, 1995). A copy of IS was introduced into the *recA*^−^ strain by lambda red recombination with pKD46_tetR. IS accumulation was performed with nine rounds of FACS-based IS accumulation.

The genotype of the cells was identified by long-read sequencing as described in a later section but with a low read depth just to identify the presence of ISs. Among the multiple strains obtained, we chose the strains that seemed to have accumulated ISs based on the sequencing results. We excluded strains that ended up amplifying ISs through tandem duplication, possibly allowing the strains to increase *rfp* while avoiding transposition-induced growth defects (example in Supplementary Fig. 2).

### Mutation accumulation experiment

The IS-accumulated strains were subjected to a mutation accumulation (MA) experiment in a nutrient-rich medium with aTc induction. For each passage, fresh LB agar plates were prepared by spreading aTc mixed in 100 µL of sterile water on preprepared LB agar plates containing 12.5 µg/mL chloramphenicol, resulting in a final aTc concentration of 3 nM. We employed a low induction concentration to avoid growth defects due to high IS activity. Chloramphenicol was added to avoid contamination of other bacteria.

Single visible colonies were randomly selected for each lineage and streaked onto a new plate to obtain isolated colonies. The plate was incubated at 32 °C for three/four overnights, with shading to avoid the degradation of aTc (two passages per week). Colonies with diameters between 2 mm and 3 mm were considered as 2.5 mm colonies and other colonies with atypical sizes were measured and recorded to infer the number of generations. The average number of generation times based on colony sizes were 216 and 540 generations after eight and twenty passages, respectively (Maeda et al., 2021).

### Genome sequencing

We used long-read sequencing to identify the genome sequences of the evolved strains. Genomes were sequenced at three points: before MA, after eight passages, and after twenty passages.

Raw reads were prepared as follows. Genome DNA was extracted from *E. coli* grown overnight using the standard phenol-chloroform method using the reagents of the Promega Wizard HMW DNA extraction kit and Phenol/Chloroform/Isoamyl alcohol (25:24:1). The extracted DNA was read using the Rapid Barcoding Kit 96 and sequenced using R.9.4.1 minion and flongle flowcells or using Native Barcoding Kit 24 V1 with R10.4.1 minion flow cells. We used v5.1.0–v5.7.5 of the MinKNOW software to run the experiments. Fastq files containing the DNA sequences were generated by ONT Guppy (v6.1.5–v6.5.7) super-accurate mode.

The reads containing sequences from individual DNA molecules were assembled to draft genomes as follows. Reads shorter than 1000 bp were filtered out using Filtlong (v0.2.0) and the remaining reads were assembled with Flye (v2.9) (Kolmogorov et al., 2019). The median N50 of post-filter reads was 22.3 kbp, and the median read depth was ×30.3 (Supplementary Table 4). Some genomes failed to assemble automatically into a single circular contig using Flye due to the presence of repeats longer than 100 kbp (e.g., L02-4 passage 20, Fig. 3.A). Such genomes were manually resolved by visualizing the assembly graph using Bandage (v0.9.0) based on the network of contigs and read depths (Wick et al., 2015). When resolving these repeats, we chose the organization resulting in the most parsimonious number of rearrangement events. Due to this limitation, inversions between the repeats could have been missed, and we might have underestimated their numbers.

The draft genomes were refined before the analysis of structural variations (SV). Assembled draft genomes were polished using medaka (v1.6.0) as follows. Fastq reads were mapped to the draft genome using minimap2 (v2.24) (Li, 2018). Variants were called using medaka consensus and medaka variant. Polished sequences were generated using bcftools (v1.15) (Danecek et al., 2021) consensus command, applying the called variants in VCF format to the draft genome. Note that we did not polish the genomes by short-read sequences, and the analyzed genomes contain false point mutations and small indels. Nevertheless, the impact of these errors on SV analysis appears negligible, as evidenced by the general consistency of results among descendants of the same ancestor and the contiguity of the assembled genomes excluding those with repeats over 100 kbp.

### Identification of IS insertion sites

To detect ISs even if they have altered structures as in Fig. 5, we adopted the following procedure. Sequences significantly (e-value < 1*e* − 10) matching the IS were identified by blastn (Camacho et al., 2009) (v2.14.0+), using the sequence of IS*1*-YK2X8 as the query. The blastn hits of candidate IS sequences were clustered, permitting a gap of up to 20 bp. These hits were assumed to be parts of the same IS. Among the IS candidate sequence clusters, those that contained at least 300 bp of continuous IS matches were classified as ISs. The direction of IS was assigned based on the largest match of IS in the cluster.

We then analyzed the “IS insertion site,” the position of IS insertion based on the coordinates of IS-excluded genome sequences. We analyzed such sites to connect ISs detected in descendant genomes to those in previous sequencing rounds, even under the presence of structural mutations. For every IS detected in the descendant genome (e.g. post-twentieth passage), we identified the IS insertion sites in the genome of the predecessor (e.g. post-eighth passage), the ancestor (pre-MA), and MDS42 (the common ancestor). Sequences of ISs, including the 100 bp sequences flanking the IS in the genome of the descendants, were searched in predecessor genomes by blastn. This will generally give two loci for each IS, one for each end because IS*1* typically forms 8 or 9 bp target site duplications upon insertion (Siguier et al., 2014). If there were duplications or inversions between ISs, the loci of the two ends would not be found near each other. To identify unique insertion sites in the coordinates of the predecessor genomes, we clustered the blastn hits of all ends of ISs in both the predecessor and descendant genomes, permitting a gap of up to 10 bp by agglomerative clustering (single linkage, scikit-learn (Pedregosa et al., 2011), v0.24.2), and chose the center of the cluster as the IS insertion site. We allowed up to 10 bp gaps to allow loci separated by the target site duplications to be clustered together.

The detection of IS insertion sites revealed some errors in the assembled sequences, which were manually corrected if necessary to avoid over-counting IS-related events. Several genomes originating from the same parental strain exhibited identical structural mutations related to ISs, perhaps due to either heterogeneity within the parental strain or mutation events between sequencing the ancestor strain and the start of MA. To prevent the over-representation of IS-related events, we manually incorporated those identical mutations into the pre-MA genomes (Supplementary Table 3). Besides, the post-MA genomes of L07 and L08 had identical IS insertion sites, likely due to cross-contamination before MA. Also, the genome of L04-1 after the twenty passages was highly diverged from the genome at the eighth passage. While the genomes of the two time-points were similar compared to genomes from other lines, many ISs inserted in the eighth passage were lost in the twentieth. Hence, we excluded the comparison between the eighth and twentieth passages of L04-1 from the analysis of IS insertion sites. Thus, the per-line event counts of SVs shown in Table 1 likely show slightly conservative estimates.

### Visualization of structural variations

Genome structure changes were visualized using SyRI (v1.6.3) (Goel et al., 2019) and plotsr (v1.1.1) (Goel and Schneeberger, 2022) (Fig. 3.A, Extended Data Figs. 3–5). To circumvent the difficulty of identifying SVs in the presence of repetitive ISs, we removed the ISs from the genome sequences before running SyRI. Genomes of consecutive rounds of sequencing were aligned by minimap2. Since we did not polish the assemblies by short-read sequences, the preset `asm20` was used to accept mismatches due to small errors in the assembled genomes. Further options `-H -f 100 -r1k,10k --rmq=no` was used for compatibility with SyRI. SyRI was run using the output bam file with options `--allow-offset 20 --cigar --no-qc`. We considered that an SV had a corresponding IS if at least one of its ends was within 20 bp of an IS. The identified coordinates of the boundaries of SVs were transformed back to the original sequences containing IS using a custom script. We used the SV annotation by SyRI for inversions, but for deletions and duplications, we used the copy number changes, as explained in *Detection of deletions and duplications*.

Rearrangements of at least 100 bp were visualized using plotsr. We preprocessed the output of SyRI to make it compatible with plotsr using a custom script as follows. We reassigned the genomes in the configuration file for plotsr into chromosomes and the generations to genomes to visualize the rearrangements in one plot. Blastn matches of ISs longer than 300 bp were exported as BED files with a format compatible with plotsr. As plotsr did not display rearrangements within a synteny block, we used a custom script to reveal the nested rearrangements inside synteny blocks. Also, we reclassified CPG (Copy Gain) or CPL (Copy Loss) events into duplications or deletion, respectively, as they were also missed by plotsr. Some mutations were classified as highly diversified regions (HDRs) by SyRI and were excluded from the visualization. This includes the complex rearrangement depicted in Fig. 3 as can be seen from the lack of bands connecting the regions in Extended Data Fig. 4.

Analysis by SyRI and plotsr revealed some errors in the assembled genomes. Among regions identified as HDRs, we manually omitted the regions that were contaminated by low-complexity repeats (Supplementary Table 4). We reran the above analyses if any modification was made to the genomes.

### Analysis of IS insertion site distribution

Hotspots of IS insertion were detected using a sliding window approach. The window was set to 100 kbp with a step size of 1 kbp. The significance of the IS insertion frequency in each region was determined using binomial tests.

We discriminated simple insertions from complex insertions entailing recombination, such as ISs that underwent intrachromosomal transposition via the cointegration pathway (Biel and Berg, 1984). To do this, IS insertion sites were categorized based on the presence of the sites across generations: ‘original’ (present in both ancestor and descendant genomes), ‘lost’ (present only in the ancestor), and ‘new’ (present only in the descendant). Insertion sites were matched between genomes of different generations based on the coordinates of the predecessor’s genome using agglomerative clustering (single linkage, scikit-learn). Simple insertions were identified when an IS has ‘new’ ends within 10 bp of each other in the predecessor’s genome. This criterion is also applied to identify eight simple insertions of composite transposons. We did not include them in the simple insertions in the hotspot analyses but included them when analyzing the “local hopping” of ISs to increase the number of simple insertion events per line.

Following a previous analysis of MA of wild-type *E. coli* (Lee et al., 2016), we analyzed the distance distribution of newly inserted ISs relative to preexisting ones. This involved comparing the distribution of distances from the newly inserted ISs to their nearest preexisting ISs with randomly generated distance distributions. The random distributions were generated from 10 000 simulated insertion events per genome in pre-MA and post-eight passage genomes.

### Detection of deletions and duplications

To identify deletions and duplications, we analyzed the copy number changes. First, we identified the copy number changes based on the MDS42 genome sequence (Extended Data Fig. 2). Genome sequences were mapped against the sequence of MDS42 but with the IS inserted between *gpmA* and *galM*, simulating the post-lambda red recombination state. We used minimap2 to align the pairs of genomes and computed the depths with the depth command of pysam (Danecek et al., 2021) (v0.21.0). Small variations in copy numbers, likely due to sequencing errors, were omitted by smoothing the depth as follows. Sequences were grouped by consecutive per-base depth estimates. Then, the read depth for small groups under 20 bp was adjusted as follows. To get a conservative estimate, we first assumed a copy number of one for the small groups. If the small groups failed to integrate with larger neighboring groups after this step, they were merged with the preceding group. This will give ranges of sequences larger than 20 bp with certain copy numbers.

To identify the numbers and size distribution of deletions and duplications, the same analysis was performed but by comparing the genome sequences from two consecutive genome sequencing rounds. We assumed that duplications occurred with the above-identified groups as units. The number of duplication events to achieve the observed copy number changes was assumed to be the copy number minus one. These assumptions can potentially lead to overcounting of duplications if nested or adjacent regions were duplicated, but no group seemed to be affected, checking by manual inspection. To avoid the distribution of duplications being distorted by tandem duplications with more than two copies, each duplicated group is counted only once in Fig. 3.C regardless of the depth. In contrast, for counting the number of duplications as in Table 1, we counted the numbers including multiple duplications of the same region. Composite transpositions were identified by manual inspection of the duplicated regions.

### Identification of essential genes

To compare the genome size changes of our study with those of the LTEE (Tenaillon et al., 2016), we adjusted the genome sizes by the essential gene contents. We identified the essential gene contents of the LTEE as follows. The criteria of essentiality used in a previous study of the LTEE (Couce et al., 2017) was not comparable with the criteria of essentiality used in other strains (Baba et al., 2006; Kato and Hashimoto, 2007). Here, we assumed that essential genes of the standard K12 strain MG1655 are essential in both MDS42 and REL606. We confirmed that the genes were found in MDS42 by matching the bnumbers and names. For REL606, all genes were found by matching the bnumber or the gene names, or by manually finding the remaining orthologs by blastn.

Essential genes of our evolved strains were identified by mapping the genes of MDS42 to the evolved genomes using blastn with the default setting. We assumed the match was valid if the e-value was less than 1e–10 and the match spanned at least 95% of the gene or the match had at most 5 bp not matched. If multiple copies of an essential gene were detected in a genome, we assumed that the gene was not essential anymore. We assumed that these essential genes were the only essential sequences in the genomes to calculate the total length of essential sequences in each genome.

### Estimation of the expected length of deletions

We estimated the expected lengths of deletions based on the positions of essential genes and ISs in the genomes. Our analysis assumed that: 1) Deletions start at the ends of IS elements. 2) Deletions can extend up to, but not include, the nearest essential gene. 3) Non-IS ends of deletions are uniformly distributed across non-essential regions.

We labeled the ends of the ISs in a genome, distinguishing between the two ends of the same IS. For the *i*-th end, let *d*_*i*_ denote the distance to the nearest essential gene in the direction of that end. This distance represents the maximum possible length of a deletion originating from that specific end of an IS and also reflects the likelihood of survival following such deletion as deletions that reach essential genes are lethal.

We first assigned weights to each IS end proportional to *d*_*i*_. These weights were normalized by the sum of the distances to the closest essential genes across each sequenced genome. For genomes pre-MA and post-eight passages, the weights were adjusted by multiplying them by the number of passages to the next sequencing round, which were eight and twelve respectively, reflecting the assumption that deletions occur at a constant rate. The expected median deletion size was calculated using the normalized weights, as these weights determine the relative probability of deletion sizes. Bootstrap confidence intervals were calculated by resampling the set of IS ends.

### Detection of IS loss

We identified IS loss by comparing the genomes of two consecutive rounds of sequencing. We did a mapping basically the reverse of the analysis of IS insertion sites. 3000 bp sequences flanking each side of a copy of IS in the predecessor genome (e.g. post-eighth passage) were searched in the descendant genome (e.g. post-twentieth passage) by blastn. The hits with the lowest e-value were considered for each side of the IS. An IS in the predecessor genome was annotated as lost if the matched end positions of the flanks had a gap smaller than 100 bp (much smaller than ISs) and the two matches were on the same strand.

Note that the detection of IS loss by this criteria might still be overestimated. In L09-4, an IS-mediated deletion was detected with a copy of IS remaining at the locus. This deletion should not revert to the original sequence even if the IS was lost, yet after the twentieth passage, the IS was not detected nor was the deletion. This suggests that some detected IS losses might be due to heterogeneity within isolated colonies.

### Detection of IS variants

IS variants shown in Fig. 5 were identified by classifying all detected ISs according to size using 100 bp bins by hierarchical clustering connecting farthest neighbors (complete linkage, scikit-learn). The first IS detected with the most common length within each bin was chosen as representative of IS variants of that size range. Genes within ISs were annotated by blastn against the IS sequences with options `-evalue 1e-5 -max_target_seqs 100000`.

### Statistical analyses

Paired t-tests, Wilcoxon rank-sum test, and correlations were calculated using R stats (v4.3.1). ANOVA was performed using linear regression using the `lm` function of R stats and the Anova function from the R car package (v3.1.2). We did not include the interaction terms. Linear regression models were performed using the lm function of R stats. Most models included an intercept, except for the analysis of the correlation between IS insertion count and genome size changes. In this case, we omitted the intercept because zero IS insertions imply no evolution, and consequently, no change in genome size.

We stated that the number of insertion sites in the recA^+^ condition was comparable to nine years of evolution in Foster’s group’s MA. This was based on the observation that 111 passages were required to reach 3080 generations in their study (Lee et al., 2012) and the assumption that one passage was performed every weekday.

Binomial tests were performed using the `binom.test` function of R stats. Sliding window analyses were performed using a custom script using `binom.test`. Results were adjusted for multiple test comparisons using the Benjamini-Hochberg method via the `p.adjust` function in the R stats package. For assessing the significance of the correlation between the frequency of essential genes and insertions within regions, we performed the same analysis but against non-overlapping windows of 100 kbp.

K-sample Anderson-Darling tests were used to compare the observed distance distribution against distances from 10 000 randomly generated insertion events using `KSampleADTest` of the HypothesisTests package (v0.11.0) in Julia. From the obtained P-values, Fisher’s combined probability tests were performed using the `sumlog` function from the R metap package (v1.1).

Bootstrap confidence intervals were calculated using the `boot` function of boot (v1.3.28) in R and `bootstrap` function of scipy (v1.9.3) in Python with 1000 bootstrap samples using the bias-corrected and accelerated (BCa) method.

## Use of Large Language Models

The text has been edited with the assistance of language models, ChatGPT, Claude, and Github Copilot.

## Data availability

The raw sequencing data and the assembled genomes will be available at DDBJ upon publication.

## Code availability

The custom scripts used for the analysis will be available on GitHub upon publication (https://github.com/Yuki-Kanai/2024_IS_Expansion_MA).

## Acknowledgments

This work was supported by the Japan Society for the Promotion of Science [21J20693 to Y.K., 18H02427 to S.T., 17H06389 to C.F, 19H05626 to C.F]; and the Japan Science and Technology Agency [JPMJER1902 to C.F].

## Author contributions

Conceptualization, Y.K., S.T., and C.F.; Methodology, Y.K., A.S., S.T., and C.F.; Software, Y.K.; Formal analysis, Y.K.; Investigation, Y.K. and N.Y; Resources, A.S., S.T. and C.F.; Data curation, Y.K.; Writing—original draft, Y.K.; Writing—review & editing, Y.K., S.T., and C.F.; Visualization, Y.K.; Supervision, S.T. and C.F.; Funding acquisition, Y.K., S.T. and C.F.

## Competing interests

The authors declare no competing interests.

### Correspondence and requests for materials

Should be addressed to S.T. or C.F.

**Extended Data Fig. 1.**
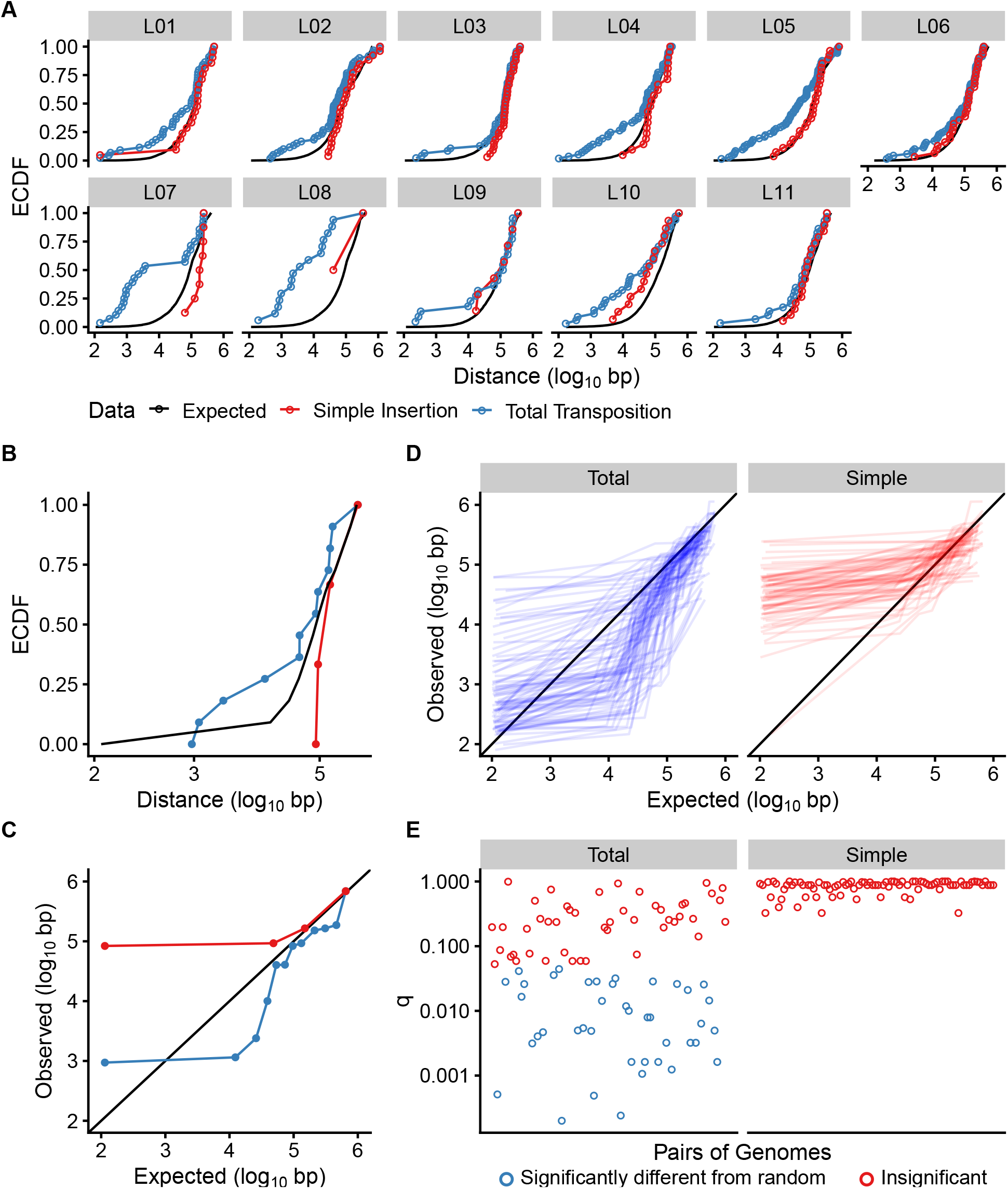
ISs were inserted closer to preexisting ISs than expected from random. The empirical cumulative distribution functions (ECDFs) of the distances between the newly inserted ISs and the ISs in the ancestor genomes were compared against their corresponding random distributions of distances. Shifts to the left indicate ISs were inserted closer to preexisting ISs than expected from random. Each point represents an additional IS insertion. This analysis follows Figure 2 of the analysis of IS insertions in *E. coli* MA by Foster’s group (Lee et al., 2016). **(B, C)** An example of a conversion from a cumulative distribution function (B, as in A) to a quantile-quantile plot (C, as in D). Colors as in A. **(D)** The quantile-quantile plot of the expected and observed distances between a newly inserted IS and an IS in the ancestor genome for 44 × 2 pairs of genomes. Shifts to the bottom right indicate local hopping of the ISs. **(E)** The q-values of the K-sample Anderson-Darling tests after correcting for multiple comparisons by the Benjamini-Hochberg method. Each point represents the divergence of a line in subfigure D from the diagonal black line. Values of all 88 pairs are shown. The Fisher’s combined probability tests in the main text used these q-values, excluding that of the twentieth passage of L04-1 due to unreliable comparison with the eighth passage genome caused by sequence divergence.

**Extended Data Fig. 2.**
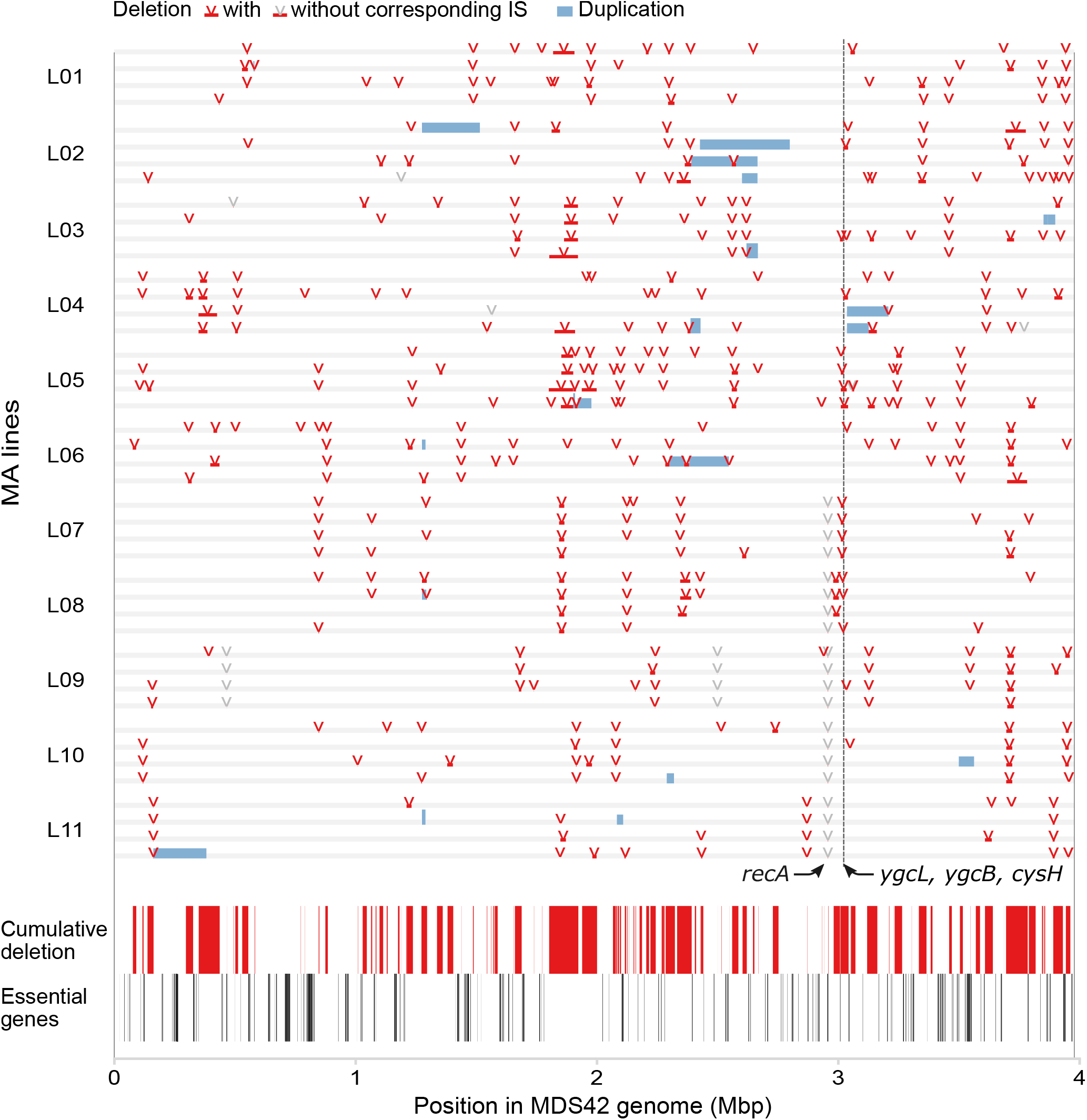
Regions of copy number variation after twenty passages of MA across 44 evolutionary lines. Lines are ordered so that L01-1 is at the top and L11-4 is at the bottom. Coordinates are based on the MDS42+IS*1* genome starting from its origin of replication. Heights indicate the copy number, where heights three times than the grey baseline indicate a copy number of three or more. *recA* was deleted before the MA in the *recA*^−^ lines. *ygcL, ygcB*, and *cysH* are the most frequently deleted genes, adjacent to each other and independently deleted in 9, 11, and 10 lines, respectively. Cumulative deleted areas of the 44 evolutionary lines and positions of essential genes according to the PEC database are shown at the bottom in red and black, respectively.

**Extended Data Fig. 3.**
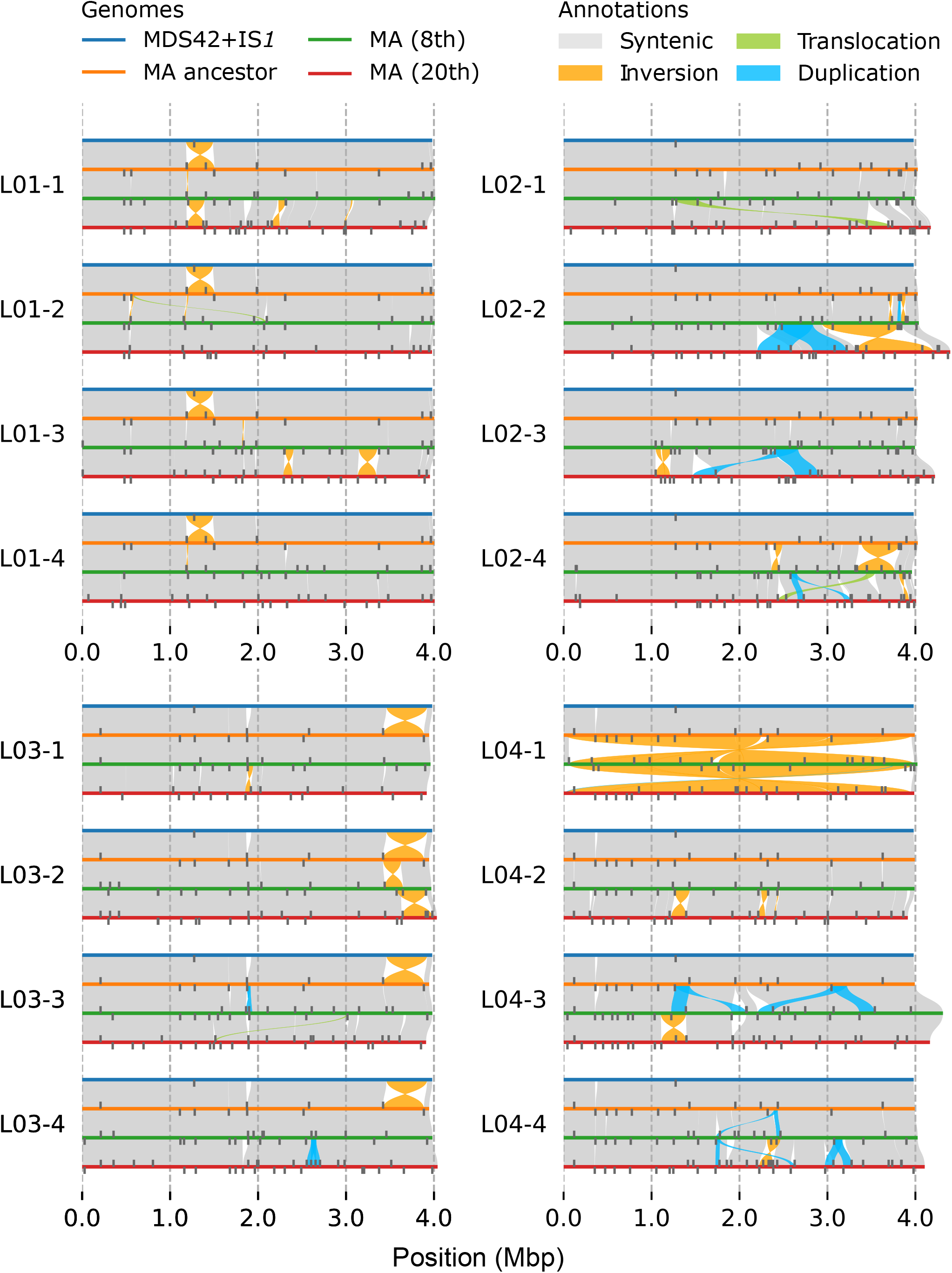
Overview of genome evolution. (L01–L04). Syntenic regions are connected by bands. Line IDs are named so that L01-2 denotes the second lineage from FACS-derived ancestor number 1.

**Extended Data Fig. 4.**
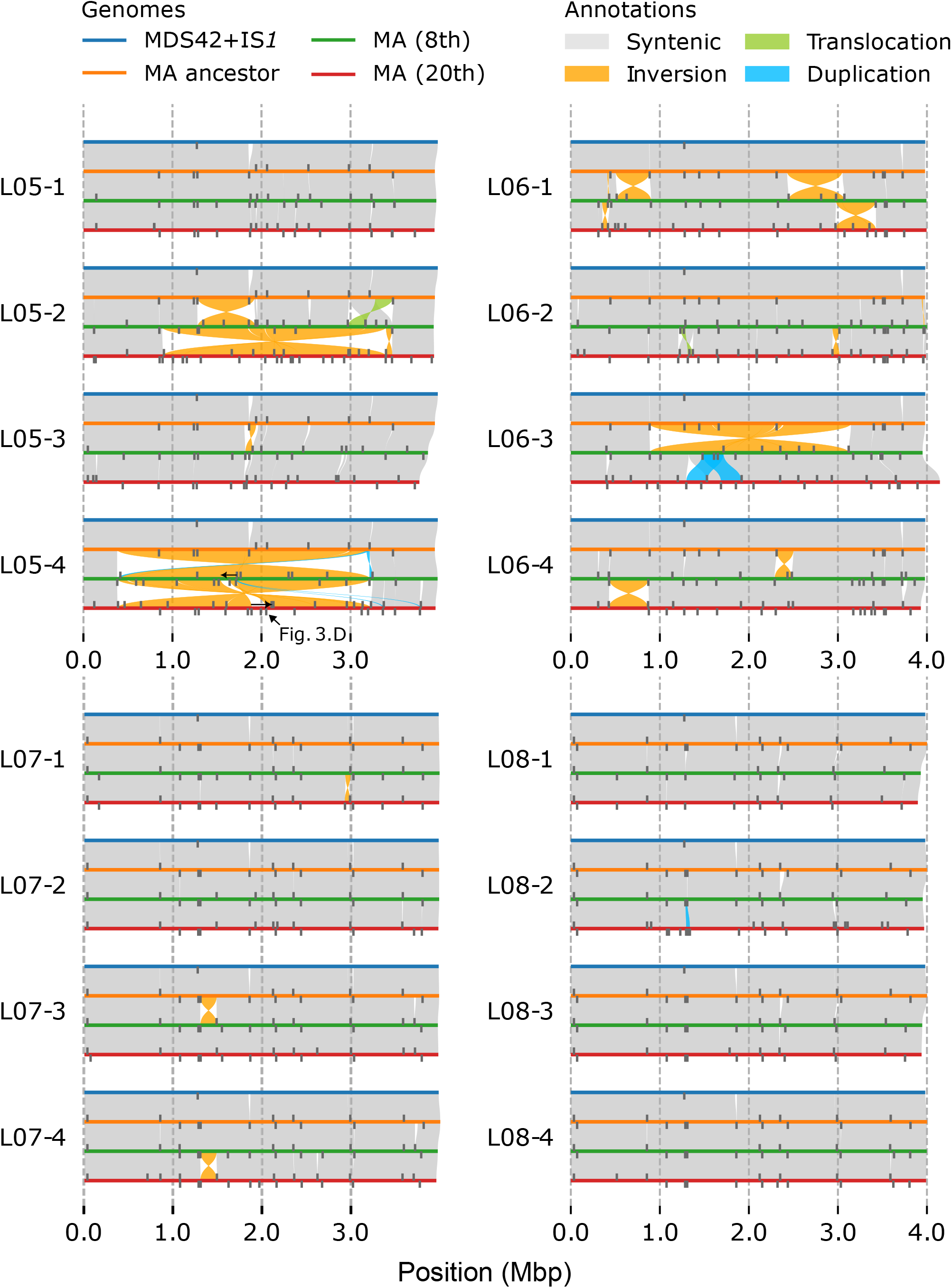
Overview of genome evolution. (L05–L08). Syntenic regions are connected by bands. Line IDs are named so that L01-2 denotes the second lineage from FACS-derived ancestor number 1. The sequences with arrows in L05-4 correspond to the complex rearrangements depicted in Fig. 3.D (arrows from left to right of the schematic figure).

**Extended Data Fig. 5.**
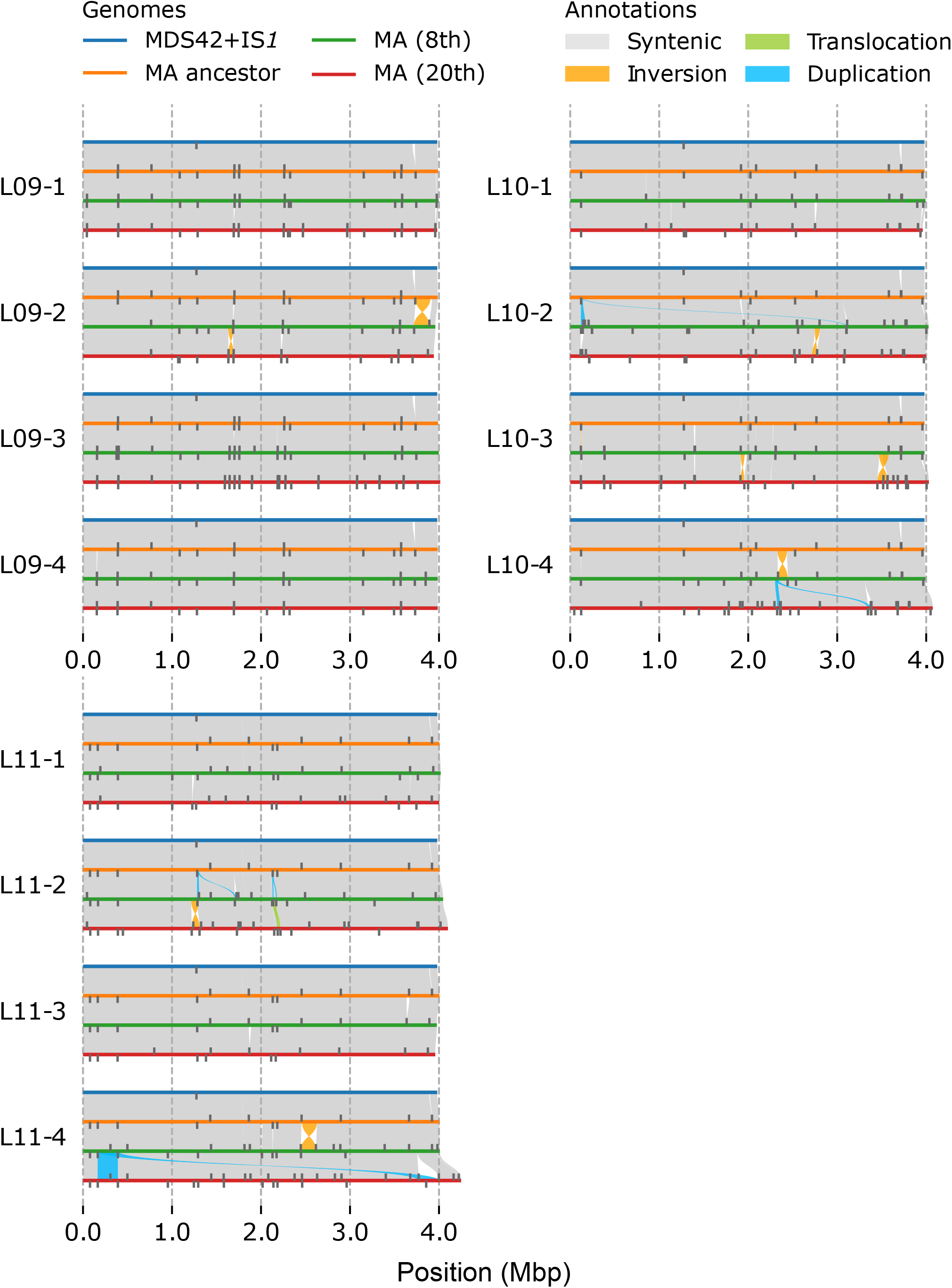
Overview of genome evolution. (L09–L11). Syntenic regions are connected by bands. Line IDs are named so that L01-2 denotes the second lineage from FACS-derived ancestor number 1.

## Supplementary Information

### Supplementary Figures

**Supplementary Fig. 1.**
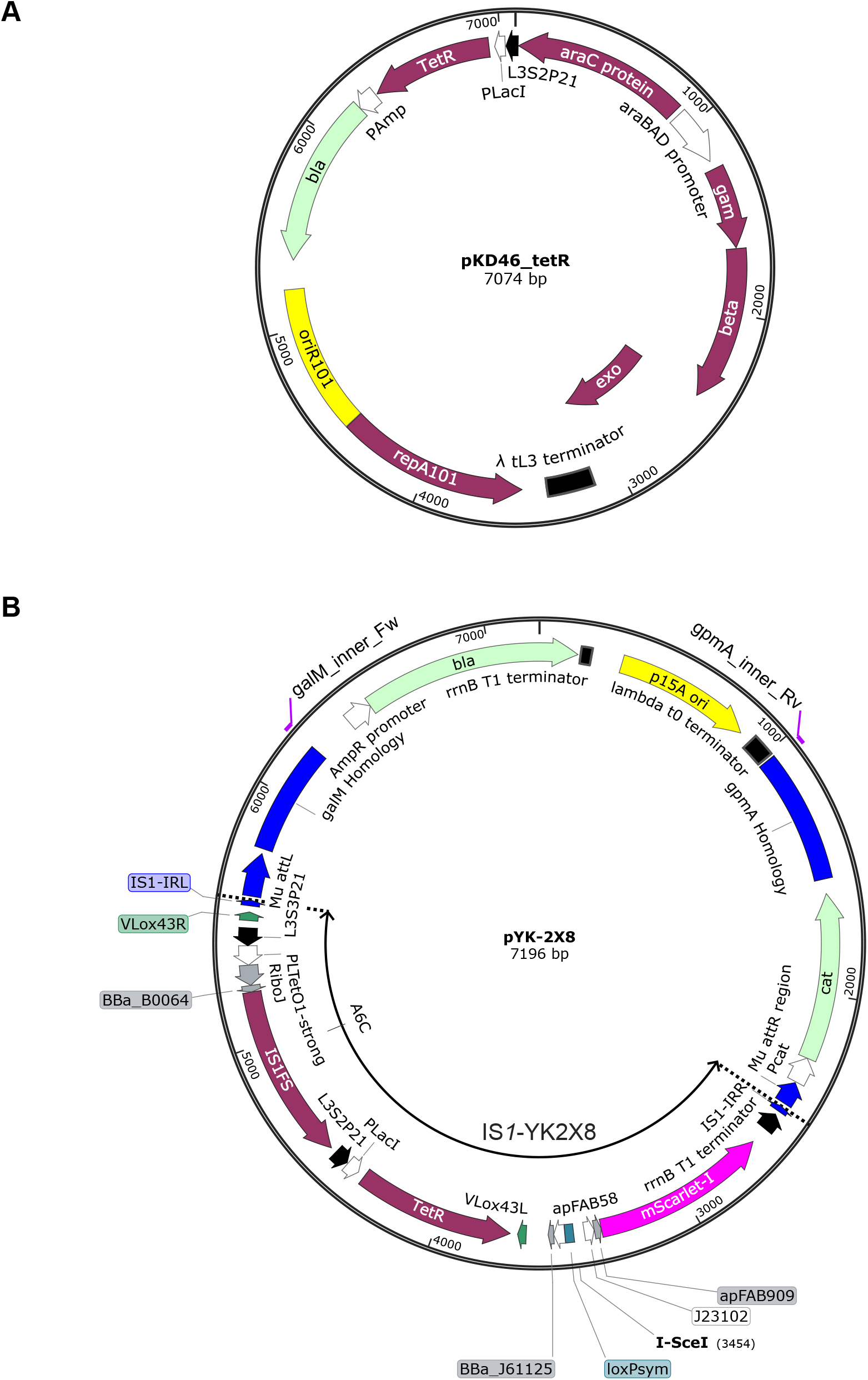
Map of plasmids constructed in this study. **(A)** pKD46_tetR. **(B)** pYK-2X8.

**Supplementary Fig. 2.**
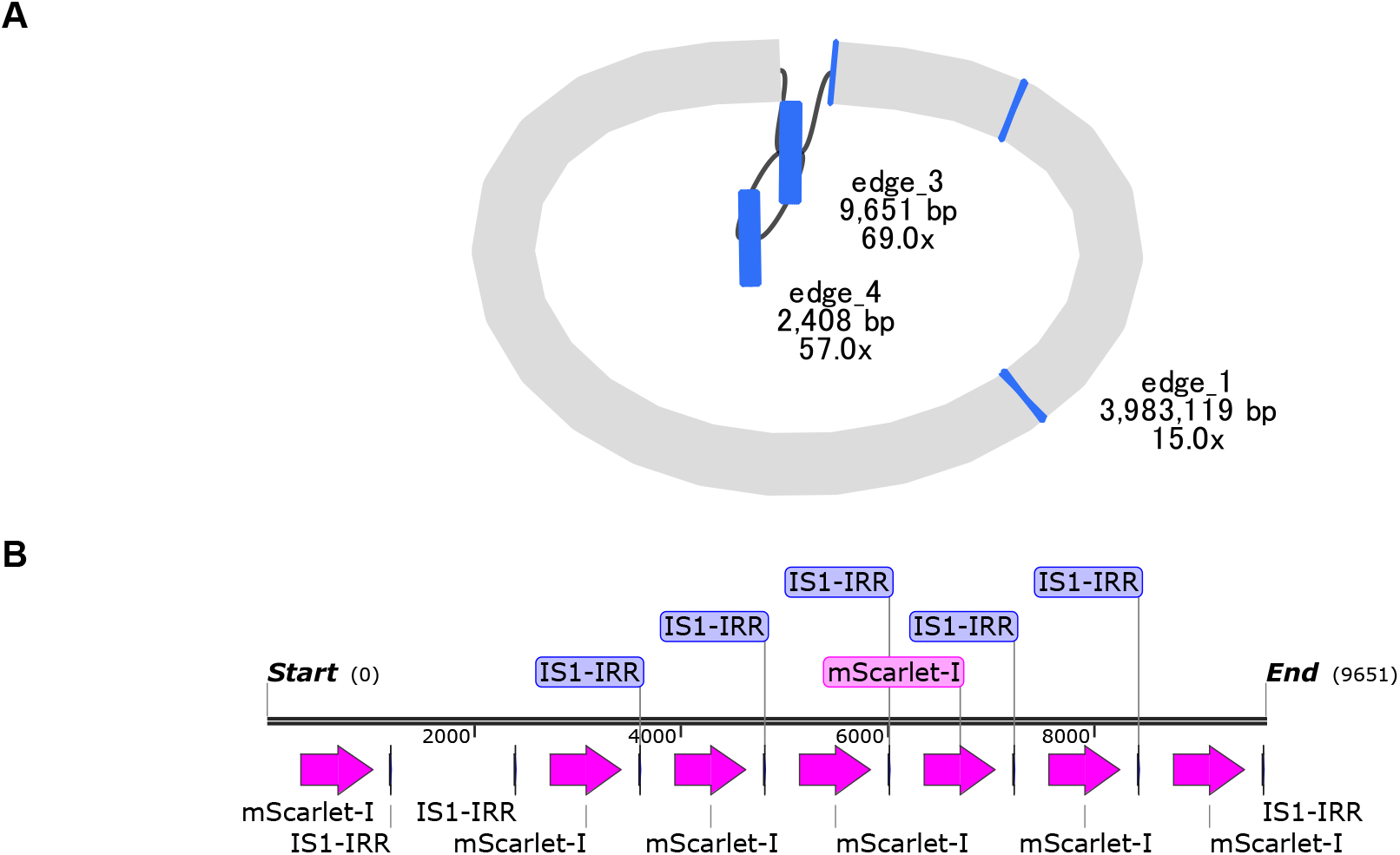
Illustration of a tandem duplication following IS enrichment via FACS. **(A)** Contig graph as visualized by Bandage (Wick et al., 2015). Blue loci represent blast hits against ISs over 100 bp. Only three loci had IS insertions were detected, which were all larger than 3 kbp. The locus including edges three and four displayed tandem duplication of *rfp*, with roughly 56 copies in total. **(B)** Gene composition within edge 3. The edge contains a tandem duplication of eight copies of *rfp* (mScarlet-I). The second *rfp* copy is not shown since its sequence had a frameshift mutation. Visualization by SnapGene.

**Supplementary Fig. 3.**
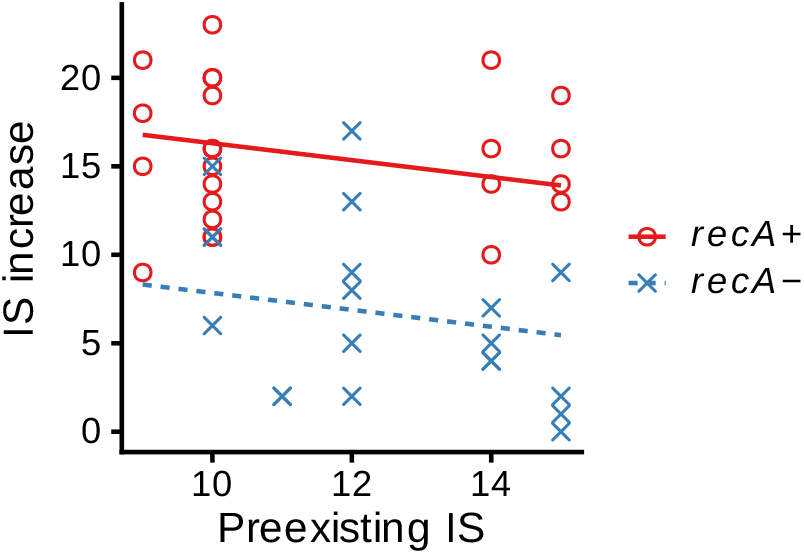
Relationship between preexisting IS count and IS copy number increase per evolutionary line. Lines represent linear regression slopes based on two factors: preexisting IS count and the presence of *recA*.

**Supplementary Fig. 4.**
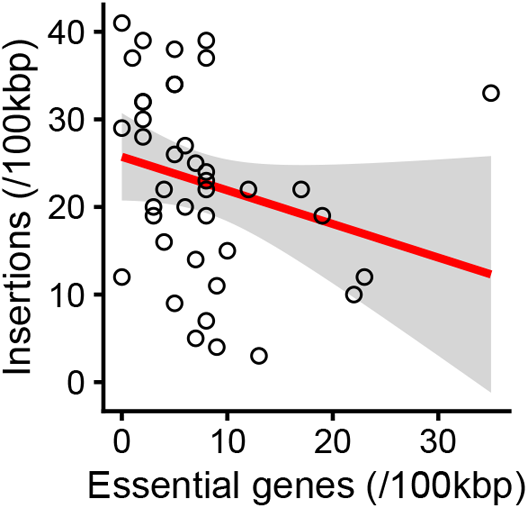
Weak anticorrelation between the number of essential genes and the number of IS insertions. The red line indicates the linear regression line with the grey area indicating the 95% confidence interval. Each point indicates a non-overlapping 100 kbp window (*n* = 40).

**Supplementary Fig. 5.**
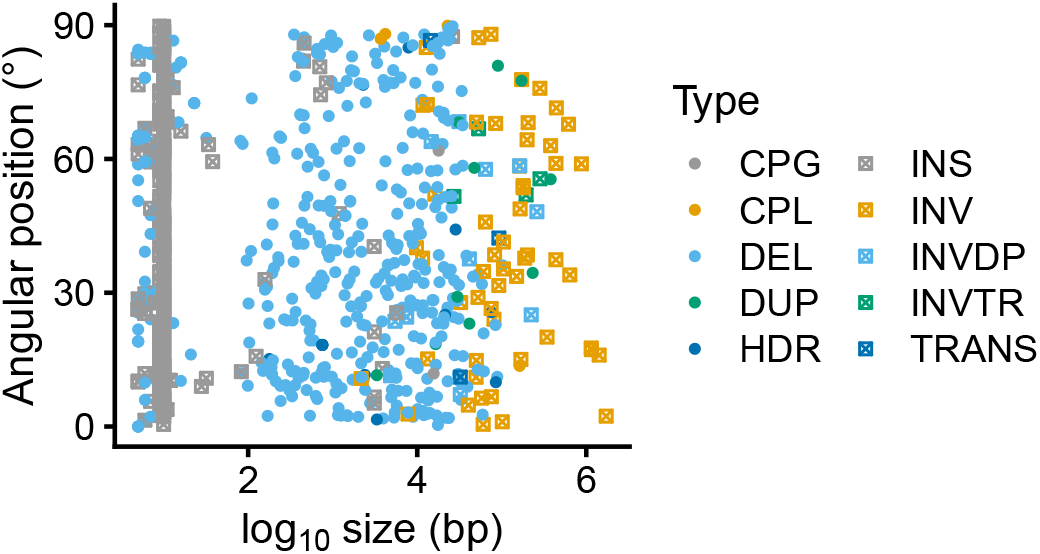
The positions and lengths of structural mutations detected by SyRI. Angular positions indicate the center of the mutations in the coordinates of their predecessor genomes. INSs clustered around 10 bp primarily represent target site duplications of IS insertions.

### Supplementary Tables

**Supplementary Table 1.**
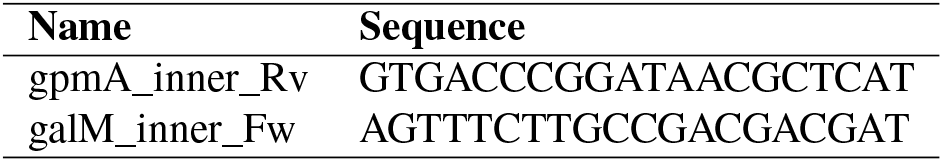
Primers used for lambda red recombination.

**Supplementary Table 2.** Composition of the M9-based rich medium.

| Substrate | Concentration | Unit | Substrate | Concentration | Unit |
| --- | --- | --- | --- | --- | --- |
| **Salts** |  |  | **Amino acids (continued)** |  |  |
| Na<sub>2</sub>HPO<sub>4</sub> | 48 | mM | Glycine | 4 | mM |
| KH<sub>2</sub>PO<sub>4</sub> | 22 | mM | Histidine | 3 | mM |
| NaCl | 8.55 | mM | Isoleucine | 3 | mM |
| NH<sub>4</sub>Cl | 9.35 | mM | Leucine | 3 | mM |
| Glucose | 22.2 | mM | Lysine | 3 | mM |
| MgSO<sub>4</sub> | 1 | mM | Methionine | 3 | mM |
| CaCl<sub>2</sub> | 0.3 | mM | Phenylalanine | 3 | mM |
|  |  |  | Proline | 3 | mM |
| **Trace elements** |  |  | Serine | 30 | mM |
| FeSO<sub>4</sub> | 10.1 | mM | Threonine | 4 | mM |
| (NH<sub>4</sub>)<sub>6</sub>Mo<sub>7</sub>O<sub>24</sub> 4H<sub>2</sub>O | 2.9 | nM | Tryptophane | 0.3 | mM |
| H<sub>3</sub>BO<sub>3</sub> | 401 | nM | Tyrosine | 0.2 | mM |
| CoCl<sub>2</sub> | 30.3 | nM | Valine | 3 | mM |
| CuSO<sub>4</sub> | 15.0 | nM |  |  |  |
| MnCl<sub>2</sub> | 80.8 | nM | **Vitamins** |  |  |
| ZnSO<sub>4</sub> | 9.7 | nM | Thiamin | 10 | μM |
|  |  |  | Riboflavin | 3 | μM |
| **Amino acids** |  |  | Nicotinamide | 100 | μM |
| Alanine | 4 | mM | Pantothenate | 100 | μM |
| Arginine | 8 | mM | Pyridoxine | 15 | μM |
| Asparagine | 3 | mM | Biotin | 1 | μM |
| Aspartate | 4.7 | mM | Folate | 0.2 | μM |
| Cysteine | 4.1 | mM | p-aminobenzoic acid | 100 | μM |
| Glutamate | 3 | mM | p-hydroxybenzoic acid | 100 | μM |
| Glutamine | 3.5 | mM | 2,3-dihydroxybenzoic acid | 100 | μM |

**Supplementary Table 3.** Rearrangements detected in multiple descendants of the same ancestor.

| Parent | Subline | Type | Start (bp) | End (bp) |
| --- | --- | --- | --- | --- |
| L01 | 1,2,3,4 | INS | 1 187 832 | 1 187 839 |
| L01 | 1,2,3,4 | INV | 1 187 832 | 1 494 557 |
| L03 | 1,2,3,4 | DEL | 1 875 064 | 1 932 897 |
| L04 | 1,2,3,4 | INS | – | 3 626 825 |
| L05 | 1,4 | DEL | 1 238 499 | 1 238 886 |
| L05 | 1,2,3,4 | INS | – | 1 916 450 |
| L05 | 1,2,3,4 | DEL | 1 862 851 | 1 916 450 |
| L05 | 1,2,3,4 | INS | 2 579 628 | – |
| L05 | 1,2,3,4 | DEL | 3 254 109 | 3 274 107 |
| L06 | 1,2,3,4 | DEL | 1 440 626 | 1 441 054 |
| L06 | 1,2,3,4 | DEL | 3 725 895 | 3 751 466 |
| L07 | 1,2,3,4 | EXC | 69 487 | 72 595 |
| L07 | 1,2,3 | DEL | 853 691 | 855 131 |
| L07 | 1,2,3 | DEL | 3 023 886 | 3 035 347 |
| L09 | 1,2,3,4 | EXC | 1 667 080 | 1 671 178 |
| L09 | 1,2,3,4 | INS | 1 701 756 | 1 701 763 |
| L09 | 1,3 | INS | 1 752 840 | 1 752 848 |
| L09 | 1,2,3,4 | DEL | 3 731 599 | 3 749 902 |

Sublines indicate the descendants of the ancestor that shared the mutation. The endpoints of some insertions are not written because they were lost due to deletion. The coordinates are based on the genomes of the ancestors before incorporating the mutations. INS: insertion, DEL: deletion, INV: inversion, EXC: excision.

**Supplementary Table 4.** Summary of the sequenced genomes.

| Sample | N50 (bp) | Depth | Contigs | Circular | Genome (bp) | Bandage | Fix |
| --- | --- | --- | --- | --- | --- | --- | --- |
| L01_Anc | 25 753 | 23.2 | 1 | Y | 4 003 851 | N | R |
| L02_Anc | 24 182 | 31.1 | 1 | Y | 4 026 846 | N | - |
| L03_Anc | 23 154 | 44.9 | 1 | Y | 3 945 915 | N | R |
| L04_Anc | 20 601 | 32 | 1 | Y | 3 992 834 | N | R |
| L05_Anc | 26 707 | 41.3 | 1 | Y | 3 947 151 | N | R |
| L06_Anc | 22 578 | 38 | 1 | Y | 3 978 605 | N | R |
| L07_Anc | 28 837 | 40.7 | 1 | Y | 3 985 195 | N | R |
| L08_Anc | 28 837 | 40.5 | 1 | Y | 4 001 201 | N | - |
| L09_Anc | 30 656 | 47.4 | 1 | Y | 3 989 727 | N | R |
| L10_Anc | 23 967 | 40.2 | 1 | Y | 3 982 402 | N | - |
| L11_Anc | 26 787 | 32.5 | 1 | Y | 4 004 368 | N | - |
| L01-1_G8 | 27 117 | 26.7 | 3 | Y | 4 014 776 | N | PC |
| L01-2_G8 | 26 011 | 18.8 | 1 | Y | 4 007 666 | N | - |
| L01-3_G8 | 28 089 | 18.4 | 1 | Y | 4 008 366 | N | - |
| L01-4_G8 | 27 400 | 42.3 | 1 | Y | 3 997 478 | N | - |
| L02-1_G8 | 24 598 | 57.8 | 2 | Y | 3 993 861 | N | P |
| L02-2_G8 | 20 008 | 215.9 | 2 | Y | 4 038 944 | N | P |
| L02-3_G8 | 28 891 | 31.3 | 1 | Y | 3 994 945 | N | - |
| L02-4_G8 | 31 124 | 30.6 | 1 | Y | 3 961 175 | N | - |
| L03-1_G8 | 16 081 | 29 | 1 | Y | 3 961 342 | N | - |
| L03-2_G8 | 17 501 | 25.6 | 1 | Y | 3 969 971 | N | - |
| L03-3_G8 | 29 572 | 51.9 | 1 | Y | 3 981 743 | N | - |
| L03-4_G8 | 19 811 | 26.9 | 1 | Y | 3 983 032 | N | - |
| L04-1_G8 | 19 140 | 21.7 | 1 | Y | 4 023 163 | N | - |
| L04-2_G8 | 21 258 | 32.8 | 1 | Y | 3 989 943 | N | - |
| L04-3_G8 | 27 148 | 24.7 | 6 | N | 4 313 004 | Y | - |
| L04-4_G8 | 23 066 | 22.3 | 1 | Y | 4 025 553 | N | - |
| L05-1_G8 | 24 561 | 18.8 | 1 | Y | 3 963 853 | N | - |
| L05-2_G8 | 17 625 | 38.9 | 1 | Y | 3 934 551 | N | - |
| L05-3_G8 | 20 665 | 12.9 | 1 | Y | 3 869 555 | N | - |
| L05-4_G8 | 18 321 | 23.4 | 1 | Y | 3 978 070 | N | - |

Supplementary Table 4 – continued from previous page
| Sample | N50 (bp) | Depth | Contigs | Circular | Genome (bp) | Bandage | Fix |
| --- | --- | --- | --- | --- | --- | --- | --- |
| L06-1_G8 | 16 610 | 18.7 | 1 | Y | 3 994 378 | N | - |
| L06-2_G8 | 15 531 | 15.4 | 1 | Y | 4 000 919 | N | - |
| L06-3_G8 | 19 241 | 19.9 | 1 | Y | 3 950 802 | N | - |
| L06-4_G8 | 16 997 | 29.6 | 1 | Y | 3 977 135 | N | - |
| L07-1_G8 | 24 886 | 18 | 1 | Y | 3 990 356 | N | - |
| L07-2_G8 | 28 732 | 22.2 | 1 | Y | 3 985 843 | N | - |
| L07-3_G8 | 21 797 | 21.3 | 1 | Y | 3 971 384 | N | - |
| L07-4_G8 | 19 128 | 40.2 | 2 | Y | 3 970 212 | N | PL |
| L08-1_G8 | 25 459 | 23.3 | 1 | Y | 3 930 436 | N | - |
| L08-2_G8 | 27 940 | 23.7 | 1 | Y | 3 952 681 | N | - |
| L08-3_G8 | 20 379 | 36.7 | 1 | Y | 3 946 890 | N | - |
| L08-4_G8 | 21 758 | 23.9 | 1 | Y | 3 997 823 | N | - |
| L09-1_G8 | 22 818 | 28.7 | 1 | Y | 3 995 870 | N | - |
| L09-2_G8 | 21 696 | 36.8 | 1 | Y | 3 958 795 | N | - |
| L09-3_G8 | 21 253 | 16.8 | 1 | Y | 3 997 607 | N | - |
| L09-4_G8 | 20 195 | 18.1 | 1 | Y | 3 985 080 | N | - |
| L10-1_G8 | 20 818 | 50 | 1 | Y | 3 992 260 | N | - |
| L10-2_G8 | 32 834 | 15.7 | 1 | Y | 4 026 576 | N | - |
| L10-3_G8 | 18 402 | 37.2 | 1 | Y | 3 981 434 | N | L |
| L10-4_G8 | 20 545 | 17.2 | 1 | Y | 3 992 207 | N | - |
| L11-1_G8 | 22 756 | 45.7 | 1 | Y | 4 019 007 | N | - |
| L11-2_G8 | 25 769 | 18.6 | 1 | Y | 4 046 183 | N | - |
| L11-3_G8 | 17 364 | 39.2 | 1 | Y | 3 975 273 | N | - |
| L11-4_G8 | 20 444 | 24.8 | 1 | Y | 4 007 862 | N | - |
| L01-1_G20 | 21 626 | 21.1 | 1 | Y | 3 923 637 | N | - |
| L01-2_G20 | 31 224 | 35.7 | 1 | Y | 3 978 375 | N | - |
| L01-3_G20 | 31 451 | 28.1 | 1 | Y | 3 954 732 | N | - |
| L01-4_G20 | 19 757 | 29.4 | 1 | Y | 4 007 510 | N | - |
| L02-1_G20 | 19 668 | 49.8 | 3 | N | 4 175 370 | Y | - |
| L02-2_G20 | 22 155 | 48.5 | 3 | N | 4 396 104 | Y | - |
| L02-3_G20 | 32 933 | 69.1 | 3 | N | 4 222 941 | Y | - |
| L02-4_G20 | 27 047 | 21.9 | 2 | N | 4 007 359 | Y | - |
| L03-1_G20 | 22 961 | 38.2 | 1 | Y | 3 918 256 | N | L |
| L03-2_G20 | 19 188 | 110 | 1 | Y | 4 033 287 | N | - |
| L03-3_G20 | 19 559 | 64.3 | 1 | Y | 3 910 841 | N | - |
| L03-4_G20 | 18 614 | 98.9 | 1 | N | 4 041 838 | Y | - |
| L04-1_G20 | 18 230 | 83.8 | 1 | Y | 3 987 308 | N | - |
| L04-2_G20 | 18 855 | 42.4 | 1 | Y | 3 914 267 | N | - |
| L04-3_G20 | 30 130 | 29.8 | 3 | N | 4 165 763 | Y | - |
| L04-4_G20 | 31 649 | 30.3 | 2 | N | 4 106 101 | Y | - |
| L05-1_G20 | 22 300 | 27 | 1 | Y | 3 943 646 | N | - |
| L05-2_G20 | 20 713 | 46.3 | 1 | Y | 3 938 244 | N | - |
| L05-3_G20 | 16 946 | 48.8 | 1 | Y | 3 773 677 | N | - |
| L05-4_G20 | 19 635 | 75.4 | 1 | Y | 3 954 835 | N | - |
| L06-1_G20 | 19 810 | 66.9 | 1 | Y | 3 995 943 | N | - |
| L06-2_G20 | 19 829 | 57.2 | 1 | Y | 3 988 727 | N | - |
| L06-3_G20 | 21 684 | 78.4 | 3 | N | 4 145 548 | Y | - |
| L06-4_G20 | 21 368 | 45.8 | 1 | Y | 3 931 227 | N | - |
| L07-1_G20 | 29 477 | 18.2 | 1 | Y | 3 986 484 | N | - |
| L07-2_G20 | 31 813 | 15.5 | 1 | Y | 3 977 066 | N | - |
| L07-3_G20 | 26 675 | 13.4 | 1 | Y | 3 976 386 | N | - |
| L07-4_G20 | 24 578 | 22.5 | 1 | Y | 3 954 512 | N | - |
| L08-1_G20 | 18 783 | 72.6 | 1 | Y | 3 897 922 | N | - |
| L08-2_G20 | 31 437 | 24 | 2 | Y | 3 969 066 | N | - |
| L08-3_G20 | 28 454 | 14 | 1 | Y | 3 939 636 | N | - |
| L08-4_G20 | 18 900 | 158.7 | 1 | Y | 3 990 215 | N | - |
| L09-1_G20 | 22 758 | 19.1 | 1 | Y | 3 980 061 | N | - |
| L09-2_G20 | 10 269 | 71.3 | 1 | Y | 3 942 000 | N | - |
| L09-3_G20 | 17 234 | 152 | 1 | Y | 4 013 673 | N | - |
| L09-4_G20 | 28 269 | 23.5 | 1 | Y | 3 984 405 | N | - |
| L10-1_G20 | 19 375 | 39 | 1 | Y | 3 960 654 | N | - |

Supplementary Table 4 – continued from previous page
| Sample | N50 (bp) | Depth | Contigs | Circular | Genome (bp) | Bandage | Fix |
| --- | --- | --- | --- | --- | --- | --- | --- |
| L10-2_G20 | 30 778 | 20.4 | 1 | Y | 3 999 127 | N | - |
| L10-3_G20 | 22 327 | 148.9 | 1 | Y | 4 030 092 | N | - |
| L10-4_G20 | 20 906 | 13.3 | 1 | Y | 4 074 199 | N | - |
| L11-1_G20 | 22 838 | 22.5 | 1 | Y | 4 002 703 | N | - |
| L11-2_G20 | 28 833 | 12.9 | 1 | Y | 4 100 257 | N | - |
| L11-3_G20 | 22 351 | 12.7 | 1 | Y | 3 958 158 | N | - |
| L11-4_G20 | 27 290 | 47.5 | 3 | N | 4 248 613 | Y | - |

N50 shows the N50 of reads after filtering short reads by filtlong. Only the data of a representative variant is shown for the ancestral genomes. The Bandage column shows whether the contigs were circularized by Bandage (Wick et al., 2015). The Fix column shows how the contigs were fixed by other means. L08-2\_G20 was not fixed by bandage despite having two contigs. The minor contig was identical to a duplicated region, indicating a copy-number variation within the population, and does not affect the conclusions of this study.
For the Fix column, the following codes are used:
R: Incorporated rearrangements from predecessors.
P: Removed plasmid contamination from other experiments.
L: Removed low complexity repeats.
C: Fixed sequence error in original IS copy due to the cross-contamination of plasmid.

